# Depth-resolved ultra-high field fMRI reveals feedback contributions to surface motion perception

**DOI:** 10.1101/653626

**Authors:** Ingo Marquardt, Peter De Weerd, Marian Schneider, Omer Faruk Gulban, Dimo Ivanov, Kâmil Uludağ

**Affiliations:** Department of Cognitive Neuroscience, Maastricht Brain Imaging Centre (MBIC), Faculty of Psychology & Neuroscience, Maastricht University, PO Box 616, 6200MD Maastricht, The Netherlands, Visiting address: Oxfordlaan 55, 6229 ER, The Netherlands; Maastricht Center of Systems Biology (MACSBIO), Faculty of Science & Engineering, Maastricht University, PO Box 616, 6200MD Maastricht, The Netherlands, Visiting address: Lenculenstraat 14, 6211 KR, Maastricht, The Netherlands; Center for Neuroscience Imaging Research Institute for Basic Science and Department of Biomedical, Engineering, N Center, Sungkyunkwan University, Seobu-ro 2066, Jangan-gu, Suwon, Korea; Techna Institute & Koerner Scientist in MR Imaging, University Health Network, 121-100 College Street, M5G 1L5 Toronto, Canada

## Abstract

Human visual surface perception has neural correlates in early visual cortex, but the extent to which feedback contributes to this activity is not well known. Feedback projections preferentially enter superficial and deep anatomical layers, while avoiding the middle layer, which provides a hypothesis for the cortical depth distribution of fMRI activity related to feedback in early visual cortex. Here, we presented human participants uniform surfaces on a dark, textured background. The grey surface in the left hemifield was either perceived as static or moving based on a manipulation in the right hemifield. Physically, the surface was identical in the left visual hemifield, so any difference in percept likely was related to feedback. Using ultra-high field fMRI, we report the first evidence for a depth distribution of activation in line with feedback during the (illusory) perception of surface motion. Our results fit with a signal re-entering in superficial depths of V1, followed by a feedforward sweep of the re-entered information through V2 and V3, as suggested by activity centred in the middle-depth levels of the latter areas. This positive modulation of the BOLD signal due to illusory surface motion was on top of a strong negative BOLD response in the cortical representation of the surface stimuli, which depended on the presence of texture in the background. Hence, the magnitude and sign of the BOLD response to the surface strongly depended on background properties, and was additionally modulated by the presence or absence of illusory motion perception in a manner compatible with feedback. In summary, the present study demonstrates the potential of depth resolved fMRI in tackling biomechanical questions on perception that so far were only within reach of invasive animal experimentation.

## Introduction

Historically, vision research has focused on the cortical response to boundaries and edges (e.g. Albrecht & Hamilton, 1982; Hubel & Wiesel, 1968). Perception, however, requires mechanisms by which areas enclosed by boundaries are ‘filled-in’. As surface perception requires spreading or integration of information over large distances, these mechanisms have been hypothesized to be localized in high-level visual areas (e.g. Dennett, 1991; Gregory, 1972; von der Heydt, Friedman, & Zhou, 2003). Several studies have indeed claimed that early visual cortex does not contribute to the processing of surfaces (Cornelissen, Wade, Vladusich, Dougherty, & Wandell, 2006; Friedman, Zhou, & von der Heydt, 2003; Perna, Tosetti, Montanaro, & Morrone, 2005). Nevertheless, a large number of human fMRI studies (Hsieh & Tse, 2010; Kok & de Lange, 2014; Mendola, Dale, Fischl, Liu, & Tootell, 1999; Pereverzeva & Murray, 2008; Sasaki & Watanabe, 2004) as well as cat (Rossi & Paradiso, 1999; Rossi, Rittenhouse, & Paradiso, 1996) and monkey electrophysiological recording studies (De Weerd, Gattass, Desimone, & Ungerleider, 1995; Komatsu, Kinoshita, & Murakami, 2000; Lamme, 1995; Lamme, Rodriguez-Rodriguez, & Spekreijse, 1999; Lu & Roe, 2007; Roe, Lu, & Hung, 2005; Zipser, Lamme, & Schiller, 1996; reviewed in Lamme & Roelfsema, 2000; and in Komatsu et al., 2000) have demonstrated retinotopic signals in response to the perception of surface brightness, colour, and texture. These surface-related neural signals in early visual cortex have raised the question to what extent they reflect feedback. As feedback projections target predominantly superficial and deep layers in early visual cortex (Anderson & Martin, 2009; Rockland & Pandya, 1979; Rockland & Virga, 1989), this leads to a clear prediction for activity distributions across cortical depth induced by feedback. In the domain of surface perception, only a handful of neurophysiological studies in animals have successfully tested layer-specific distributions of activity during feedback. Using texture-defined surfaces, two neurophysiological studies in monkeys (Self, van Kerkoerle, Supèr, & Roelfsema, 2013; van Kerkoerle et al., 2014) revealed complex temporal patterns engaging both deep and superficial layers. A single human fMRI study using a static surface induced in a Kanizsa display (Kok, Bains, van Mourik, Norris, & de Lange, 2016) reported cortical deep layer activity compatible with a role of feedback in surface perception. These experiments align with anatomical data indicating that feedback projections can target both superficial and deep layers. Recent optogenetics studies in mice have moreover confirmed that the correlates of feedback in V1 causally depend on activity in high-level visual cortex (Schnabel et al., 2018).

Surface perception is thought to interact tightly with mechanisms of contour reconstruction. A number of computational models of surface perception (Grossberg, 1987a, 1987b; see also Keil, Cristóbal, Hansen, & Neumann, 2005) have proposed that diffusion-like spreading in a surface feature system is contained within proper retinotopic bounds by local inhibition delivered by boundary representations. Neurophysiological observations of contour-related responses in V2 (von der Heydt, Peterhans, & Baumgartner, 1984) and in V1 (Grosof, Shapley, & Hawken, 1993) and surface related responses in V1, V2 and V3 (De Weerd et al., 1995; Huang & Paradiso, 2008) have emphasized the role of early visual areas in this interaction between surface and contour processing.

Separating responses to edges from responses to the interior of a surface is of utmost importance, as contour responses themselves involve feedback (Lee & Nguyen, 2001), (Wokke, Vandenbroucke, Scholte, & Lamme, 2013), and may show a depth distribution of activity in early visual cortex similar to that elicited by responses to surfaces. In the only depth-specific human fMRI study on surface perception to date, Kok et al. presented participants with Kanizsa stimuli containing illusory surfaces and contours (Kok et al., 2016). The illusory stimuli caused a response at deep cortical depths in V1, suggesting feedback originating from higher cortical areas. However, due to stimulus design and choice of the region-of-interest (ROI), the feedback related signal could be due to the illusory contour or to the illusory surface, because the ROI could have captured activity related to both.

Research using visual illusions to study the neural correlates of surface perception has predominantly used static surfaces with induced percepts of brightness, colour, or texture, while these features were physically absent in these surfaces. We are aware of only one previous study that measured responses to induced motion of a uniform surface, i.e. without local changes in retinotopic input (Akin et al., 2014). In fMRI studies focusing on motion interpolation, feedback-related responses in V1 were most likely driven by contours rather than surfaces (Meng, Remus, & Tong, 2005; Muckli, Kohler, Kriegeskorte, & Singer, 2005; Seghier et al., 2000), and other motion-related V1 responses may have been driven by local elements in a non-uniform surface (Muckli, Singer, Zanella, & Goebel, 2002).

By contrast, here we used a stimulus (adapted from Akin et al., 2014, see Supplementary Table 1 for a detailed comparison of stimulus parameters) that consisted of a centrally fixated, luminance-defined disk, of which a sector was removed. The removed sector was limited to the right hemifield, and rotated clockwise and anticlockwise within the right hemifield, thereby inducing a motion percept of the disk. In the left hemifield, the entire half of the disk was static, remained physically identical, and did not contain local elements inducing the movement percept. Two control conditions that eliminated the illusory motion kept the half of the disk in the left hemifield identical as well. That is, the three stimuli differed in global and local perceptual quality, while being physically identical in the left half of the visual field.

These stimuli, hence, provide several advantages: First, because the motion percept is induced without relying on local elements, an fMRI correlate of surface motion cannot be reduced to merely a modified processing of local elements. Second, because the retinal image of illusory and control stimuli was identical in the left hemifield, and because transcallosal connections are restricted to the vertical meridian in primate early visual cortex (Clarke & Miklossy, 1990; Essen & Zeki, 1978; Glickstein & Whitteridge, 1976; Wong-Riley, 1974), any difference between stimulus conditions can be attributed unambiguously to top-down feedback effects. Third, the stimulus was large enough so that contributions to the fMRI signal from the surface were separable from contributions from the contour, enabling any feedback signal to be attributed solely to the surface.

Furthermore, we used ultra-high field (UHF) 7T fMRI to test whether the attribution of motion to a locally static, luminance-defined surface leads to a depth-resolved pattern of activity consistent with feedback processing in early visual cortex. While the tools to perform layer-specific recordings have been available in invasive neurophysiology in animals for decades, the analysis of depth-specific activity in humans has only recently become within reach thanks to UHF fMRI and advances in data analysis (Guidi, Huber, Lampe, Gauthier, & Möller, 2016; Huber et al., 2015; Koopmans, Barth, & Norris, 2010; Koopmans, Barth, Orzada, & Norris, 2011; Marquardt, Schneider, Gulban, Ivanov, & Uludağ, 2018; Olman et al., 2012; Polimeni, Fischl, Greve, & Wald, 2010; Ress, Glover, Liu, & Wandell, 2007). Our analysis included not only V1 (as in Kok et al., 2016), but was extended to V2 and V3.

Notably, in the non-depth resolved fMRI study that inspired our stimulus design, a negative BOLD response was reported in response to the grey figure region presented on a dark background. Irrespective of whether the BOLD response to the grey figure was negative or positive, we hypothesized that the illusory perception of surface motion would be associated with enhanced activity in superficial and/or deep layers compared to control conditions, in accordance with a contribution of feedback in early visual cortex.

## Methods

### Experimental design

Healthy participants (*n*=9, age between 18 and 44 years, mean (SD) age 27.6 (7.3) years) gave informed consent before the experiment, and the study protocol was approved by the local ethics committee of the Faculty for Psychology & Neuroscience, Maastricht University. Subjects were presented three visual stimuli: The main experimental stimulus was a ‘Pac-Man’ figure rotating around its centre (Figure 1A). There were two control conditions: First, the same Pac-Man figure as in the main condition was presented statically, i.e. without rotating around its centre (Figure 1B). Second, the third stimulus consisted of a large, stationary wedge on the left side, and a smaller, rotating wedge on the right side (at the same location as the ‘mouth’ of the Pac-Man; Figure 1C). We will henceforth refer to these three conditions as ‘Pac-Man dynamic’, ‘Pac-Man static’, and ‘control dynamic’, respectively.

**Figure 1.**
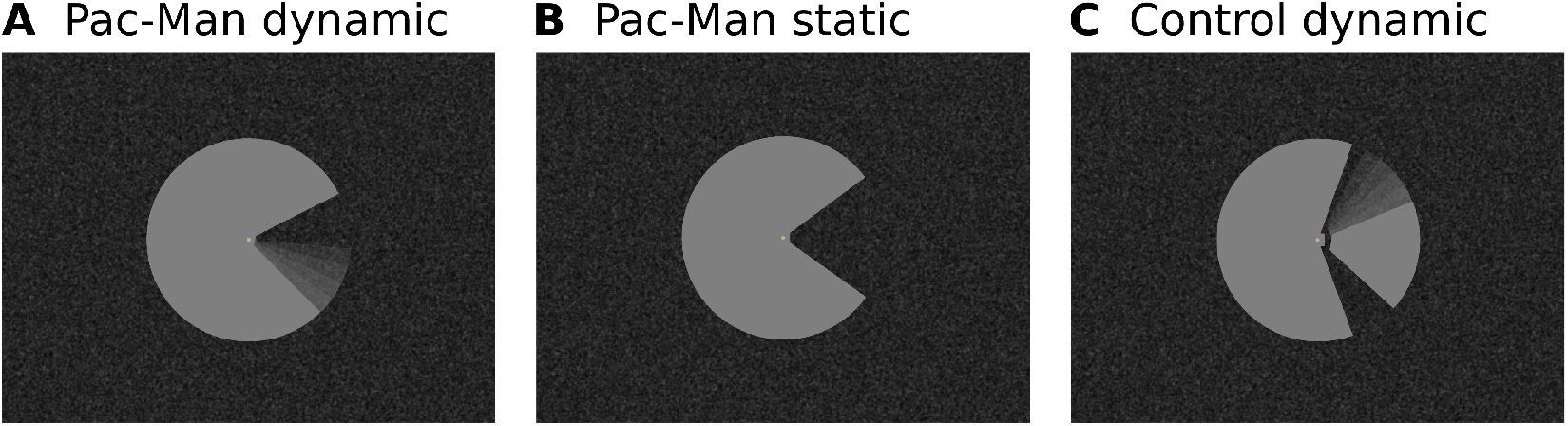
Stimulus Design. (A) A ‘Pac-Man’ figure rotating about its centre served as the main experimental stimulus. At the beginning of each stimulus block, the ‘mouth’ was centred on the horizontal meridian (i.e. mirror-symmetric about the horizontal meridian). The ‘mouth’ had a circular arc of 70° (±35° from the right horizontal meridian), and rotated clockwise and anticlockwise by ±35° (with respect to the right horizontal meridian), at a rate of 0.85 cycles per second. This experimental condition is referred to as ‘Pac-Man Dynamic’. (B) In the first of two control conditions, the same Pac-Man figure as in (A) was presented statically, i.e. without rotating about its centre. This condition is referred to as ‘Pac-Man static’. (C) In the second control condition, a figure consisting of a stationary wedge on its left side, and a smaller, rotating wedge on its right side was presented. The movement of the right-hand wedge was similar to that of the ‘mouth’ of Pac-Man dynamic; i.e. it started of centred of the horizontal meridian, and rotated with the same frequency and angular displacement as the ‘mouth’ of Pac-Man dynamic. The rotating, right-hand wedge had a circular arc of 65°, and the stationary, left-hand wedge had a circular arc of 220°. This condition is referred to as ‘control dynamic’. All three stimuli had a diameter of 7.5° visual angle. In (A) and (C), the angular position of the ‘mouth’ and the wedge were modulated sinusoidally, in order to create the impression of a smooth, natural movement. Importantly, the Pac-Man dynamic stimulus is perceived to rotate as a whole, whereas the control dynamic stimulus creates the impression of a rotating wedge on the right, and a stationary wedge on the left. At the same time, the retinal image of all three stimuli is identical in the left visual field. All stimuli were presented on a textured random noise background in order to enhance figure-ground segmentation. The stimuli, including the texture background, were adapted from Akin et al. (2014). Videos of the stimuli are available online (https://doi.org/10.5281/zenodo.2583017).

All three stimuli had a diameter of 7.5° visual angle. The ‘mouth’ of the Pac-Man had a circular arc of 70° (±35° from the right horizontal meridian). In the Pac-Man dynamic condition, the ‘mouth’ of the Pac-Man rotated clockwise and anticlockwise by ±35°, at a rate of 0.85 cycles per second. The angular position of the ‘mouth’ was modulated sinusoidally in order to create the impression of a smooth, natural movement. In the control dynamic condition, the right-hand wedge rotated with the same frequency and angular displacement as the ‘mouth’ of the Pac-Man. The rotating, right-hand wedge had a circular arc of 65°, and the stationary, left-hand wedge had a circular arc of 220°. As a result, the Pac-Man dynamic stimulus is perceived to rotate as a whole, whereas the control dynamic stimulus creates the impression of a rotating wedge on the right and a stationary wedge on the left. Importantly, the retinal image of all three stimuli is identical in the left visual field.

All stimuli were presented on a textured random noise background as was done in Akin et al.. (2014), who included the texture to increase figure ground segregation. The background texture pattern was static, and was displayed throughout each run (i.e. also during rest periods). The texture pattern was created by randomly drawing pixel intensity values from a Gaussian distribution, and filtering the resulting image with a uniform kernel (kernel size 6 x 6 pixel). Before applying the uniform filter, the random Gaussian distribution of pixel intensities had a mean of 40 units and a standard deviation of 60 units (8-bit unsigned integer RGB pixel intensities, i.e. range 0 to 255). The granularity of the texture pattern is a function of the size of the filter kernel, and of the width of the Gaussian distribution, from which the pixel intensities are drawn. The relation between pixel intensity and luminance on our projection system was given by *y* = −78.8 × *x*^3^ + 78.7 × *x*^2^ + 317.2 × *x* + 163.3, where *x* represents the pixel intensity (in Psychopy convention, i.e. range −1.0 to 1.0), and *y* corresponds to luminance (in cd/m^2^). These values are based on measurements taken with a photometer (Konica Minolta CS-100A), and subsequent least-squares fitting of several functions, of which a third-degree polynomial provided by far the best fit. The mean luminance of the texture background was 8 cd/m^2^, and the experimental stimuli (‘Pac-Man’ and control stimulus) had a uniform luminance of 163 cd/m^2^. Videos of the stimuli are available online (https://doi.org/10.5281/zenodo.2583017).

Stimuli were created with Psychopy (Peirce, 2007, 2008) and projected onto a translucent screen mounted behind the MRI head coil, via a mirror mounted at the end of the scanner bore. The three stimulus conditions were presented in separate runs and in random order (see Supplementary Figure 1). Stimuli were presented in a block design with block durations of 10.4 s and variable rest periods in random order (18.7 s, 20.8 s, or 22.9 s). Each run began with an initial rest period with a fixed duration of 20.8 s, and ended with a rest period of one of the three possible durations. All lights in the scanner room were switched off during the experiment, and black cardboard was placed on the inside of the MRI transmit coil in order to minimise light reflection. Each subject completed six functional runs (two for each stimulus condition; with the exception of one subject, who completed three repetitions each of the Pac-Man dynamic and control dynamic conditions, and two for Pac-Man static). The total duration of a run was 520 s.

Participants were asked to fixate a central dot throughout the experiment and to report pseudo-randomly occurring changes in the dot’s colour by button press. These targets were presented for 800 ms, with a mean inter-trial interval of 30 s (range ±10 s). No targets appeared during the first and last 15 s of each run. The timing of the colour changes was arranged such that the predicted haemodynamic responses to the experimental stimulus and to the colour changes are uncorrelated. To this end, a design vector representing the stimulus blocks and a design vector containing pseudo-randomly timed target events were separately convolved with a gamma function serving as model for the haemodynamic responses. The correlation between the predicted responses to the stimulus blocks and to the target events was calculated, and if the correlation coefficient was above threshold (*r* > 0.001), a new pseudo-random design matrix of target events was created. This procedure was repeated until the correlation was below threshold, separately for each run.

In an additional run, retinotopic mapping stimuli were presented for population receptive field estimation, allowing us to delineate early visual areas V1, V2, and V3 on the cortical surface (Dumoulin & Wandell, 2008). Please see Supplementary Material for details on the stimulus design of the population receptive field mapping paradigm.

In order to determine whether the responses are sustained or transient (Horiguchi, Nakadomari, Misaki, & Wandell, 2009; Uludağ, 2008), we acquired an additional experimental run for the Pac-Man dynamic condition with longer block durations in a subset of subjects (*n*=5). The additional run had a duration of 424 s, during which the dynamic Pac-Man stimulus was presented five times for 25 s, interspersed between rest blocks of 50 s. As in the main experiment, subjects performed a central fixation task.

### Control experiment

A further control experiment was conducted to investigate the role of the stimulus shape and of the background in the processing of a surface stimulus. Two uniform surface stimuli were presented: A central disk from which a sector was removed (i.e. identical to the ‘Pac-Man static’ in the main experiment), and a central square. Both stimuli were identical in luminance and area. The square had a side length of 6.65° visual angle. Both stimuli were presented under two background conditions: either on a uniform, dark grey background, or on a random texture background (same as in the main experiment). The two background conditions (i.e. uniform/texture) were presented in separate experimental runs, whereas the two stimulus shapes (i.e. Pac-Man/square) were presented in random order within runs. Stimulus blocks had a duration of 12.4 s, and were interspersed with variable rest blocks of 22.9 s, 25.0 s, or 27.0 s. The uniform background and the random texture pattern had a luminance of 8 cd/m^2^, and the surface stimuli (Pac-Man & square) had a luminance of 163 cd/m^2^ (same as in the main experiment). The control experiment was conducted in a separate session. Two subjects completed six experimental runs each (three with uniform background, three with texture background). Videos of the stimuli are available online (https://doi.org/10.5281/zenodo.2583017). As in the main experiment, retinotopic mapping runs were acquired in the same session.

### Data acquisition & preprocessing

Functional MRI data were acquired on a 7 T scanner (Siemens Medical Systems, Erlangen, Germany) and a 32-channel phased-array head coil (Nova Medical, Wilmington, MA, USA) using a 3D gradient echo (GE) EPI sequence (TR = 2.079 s, TE = 26 ms, nominal resolution 0.8 mm isotropic, 40 slices, coronal oblique slice orientation, phase encode direction right-to-left, phase partial Fourier 6/8; Poser, Koopmans, Witzel, Wald, & Barth, 2010). We also acquired whole-brain structural T1 images using the MP2RAGE sequence (Marques et al., 2010) with 0.7 mm isotropic voxels, and a pair of five SE EPI images with opposite phase encoding for distortion correction of the functional data (TR = 4.0 s, TE 41 = ms; position, orientation, and resolution same as for the GE sequence; Feinberg et al., 2010; Moeller et al., 2010; Setsompop et al., 2012).

Motion correction was performed using SPM 12 (Friston, Williams, Howard, Frackowiak, & Turner, 1996), and the data were distortion corrected using FSL TOPUP (Andersson, Skare, & Ashburner, 2003). Standard statistical analyses were performed using FSL (Smith et al., 2004), fitting a general linear model (GLM) with separate predictors for the three stimulus conditions and a nuisance predictor for the target events of the fixation task. In order to account for both sustained and transient responses, each of the three stimulus conditions was modelled with two predictors: one based on a ‘boxcar function’ over the entire stimulus duration, and the other based on a delta function at stimulus onset and offset. (Only one predictor was used for the short target events.) All GLM predictors were convolved with a double-gamma haemodynamic response function. Highpass temporal filtering (cutoff = 35 s) was applied to the model and to the functional time series before GLM fitting. The parameter estimates obtained from the GLM were converted into percent signal change with respect to the initial pre-stimulus baseline (i.e. the first 20.8 s of each run). Population receptive field mapping (Dumoulin & Wandell, 2008) was performed using publicly available python code (https://doi.org/10.5281/zenodo.1475439) and standard scientific python packages (Numpy, Scipy, Matplotlib, Cython; Behnel et al., 2011; Hunter, 2007; Millman & Aivazis, 2011; Oliphant, 2007; van der Walt, Colbert, & Varoquaux, 2011). In order to facilitate reproducibility, the complete analysis pipeline was containerised within docker images (Halchenko & Hanke, 2012; Kaczmarzyk et al., 2017).

Cortical depth sampling requires a high level of spatial accuracy. In order to detect and remove low-quality data based on a quantifiable and reproducible exclusion criterion, we calculated the spatial correlation between each functional volume and the mean EPI image of that session after motion correction and distortion correction (see Marquardt et al., 2018, for details). If the mean correlation coefficient of the volumes in a run was below threshold (*r* < 0.95), that run would have been excluded from further analysis. However, no runs were excluded based on the spatial correlation criterion. Moreover, it was important for subjects to be awake and to maintain fixation throughout the experiment. Therefore, runs in which subjects had detected less than 70% of targets were excluded from the analysis. This led to the exclusion of all runs from one subject. All other subjects had detected more than 70% of targets on all runs (mean hit rate for all subjects = 93%, standard deviation = 18%; mean hit rate after exclusion criterion = 98%, standard deviation = 5%).

### Segmentation & cortical depth sampling

Separately for each subject, the anatomical MP2RAGE images were registered to the mean functional image. In order to avoid downsampling of the anatomical images during registration, the mean functional image of each subject was upsampled to a resolution of 0.4 mm isotropic before registration (using trilinear interpolation). Thus, during registration of the anatomical images to the upsampled mean functional image, the anatomical images were indirectly upsampled (from 0.7 mm to 0.4 mm isotropic). This upsampling of anatomical images is beneficial for fine-grained tissue type segmentation, because it allows for better separation of adjacent sulci (avoiding erroneous grey matter ‘bridges’). The anatomical images were roughly aligned in a first registration step based on normalized mutual information, followed by boundary-based registration (Greve & Fischl, 2009; Jenkinson, Bannister, Brady, & Smith, 2002; Jenkinson & Smith, 2001). The registered MP2RAGE images were used for tissue type segmentation. Initial tissue type segmentations was created with FSL FAST (Zhang, Brady, & Smith, 2001). These initial segmentations were semi-automatically improved using the Segmentator software (Gulban, Schneider, Marquardt, Haast, & De Martino, 2018) and ITK-SNAP (Yushkevich et al., 2006). These corrections of the segmentations obtained from FSL FAST were based on the T1 image from the MP2RAGE sequence, and aimed to remove mistakes in the definition of the white/grey matter boundary and at the pial surface.

The final white and grey matter definitions were used to construct cortical depth profiles using volume-preserving parcellation implemented in CBS-tools (Bazin et al., 2007; Waehnert et al., 2014). Specifically, the cortical grey matter was divided into 10 compartments, resulting in 11 depth-level images delineating the borders of these equi-volume compartments. The results from the GLM analysis, the population receptive field estimates, and event-related fMRI time courses were up-sampled to the resolution of the segmentations (i.e. 0.4 mm isotropic voxel size) using trilinear interpolation, and sampled along the previously established depth-levels using CBS-tools (Bazin et al., 2007; Waehnert et al., 2014). The depth-sampled data were projected onto a surface mesh (Tosun et al., 2004).

### ROI selection

We aimed to define ROIs in an observer-independent, quantifiable way. Only the first step of the ROI selection, i.e. the delineation of cortical areas V1, V2, and V3, was performed manually. The visual areas V1, V2, and V3 were delineated on the inflated cortical surface based on the polar angle estimates from the pRF modelling using Paraview (Ahrens, Geveci, & Law, 2005; Ayachit, 2015). Subsequently, three selection criteria were applied for each location on the cortical surface for all cortical depths (i.e. each cortical segment) contained within V1, V2, or V3. First, only segments with good population receptive field model fits were included (*R2* > 0.15, median across cortical depth levels), excluding regions that are not specifically activated (e.g. possibly due to responses to a wide range of visual angles). Second, segments with low signal intensity in the mean EPI image were excluded, in order to avoid sampling from veins and low intensity regions around the transverse sinus, which may be present due to slight imprecisions in the registration and/or segmentation. Specifically, segments with a mean EPI image intensity below 7000 at any cortical depth (i.e. minimum over cortical depths) were excluded. (The mean EPI image intensity was ∼10.000 for voxels within the brain.) Third, separate ROIs were defined for the centre of the stimulus, with eccentricities between 1° to 3° visual angle, and for the edge of the stimulus, at eccentricities between 3.5° and 4.0° visual angle (see Figure 2). The eccentricity of a segment was defined as the median eccentricity over cortical depths. The lower bound of the ROI corresponding to the stimulus centre was set to 1° (and not to 0°) in order to avoid the cortical representation of the fixation dot. Selection criteria were always applied to all cortical depths in a segment – i.e. either the entire cortical segment was included or excluded. Because the physically constant half of the stimulus was located in the left visual hemifield, the analysis was restricted to the right hemisphere (with the exception of the visual field projections, which were reconstructed from both hemispheres; Figures 5 & 6). The ROI selection described in this section, and all subsequent analysis steps were performed using standard scientific python packages (Numpy, Scipy, Matplotlib; Hunter, 2007; Millman & Aivazis, 2011; Oliphant, 2007; van der Walt et al., 2011). Percent signal change values were averaged over the ROI, separately for each cortical depth level.

**Figure 2.**
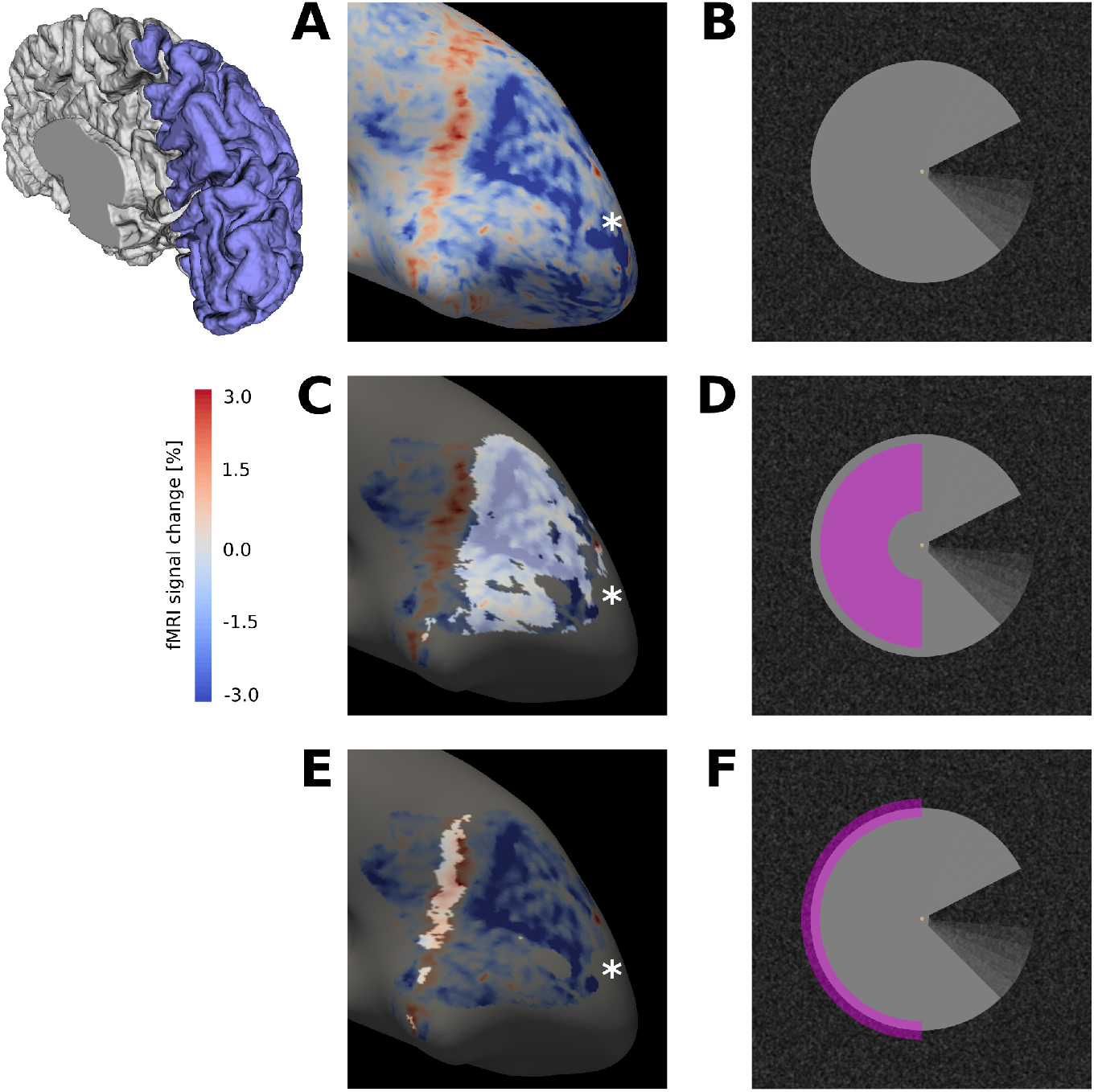
(A) Activation map for Pac-Man dynamic condition (stimulus shown in (B)), projected on the inflated cortical surface, for a representative subject (GLM parameter estimates for sustained response). An extended region of negative signal change (blue) is surrounded by a band of positive signal change (red). (C) The activation map from (A) is masked for V1, and the cortical area that retinotopically corresponds to the centre of the Pac-Man stimulus (D) is highlighted. (E) Same as (C), but the cortical area that contains the retinotopic representation of the edge of the Pac-Man stimulus (F) is highlighted. The band of positive signal change corresponds to the retinotopic representation of the edge of the Pac-Man stimulus. The areas highlighted in (C) and (E) were selected as ROIs for the stimulus centre and edge, respectively. Discontinuities in the ROIs are due to thresholding of the retinotopic map (*R2* > 0.15). The asterisk marks the approximate location of the cortical representation of the fovea (A, B, C). The schematic of a right hemisphere next to (A) indicates the approximate location of the inflated surface in (A, C, E), highlighted in blue.

### Draining effect spatial deconvolution

Cortical depth-specific fMRI using GE sequences is affected by a venous bias caused by ascending draining veins, resulting in an fMRI signal increase towards the cortical surface (Koopmans et al., 2011; Markuerkiaga, Barth, & Norris, 2016; see Uludağ & Blinder, 2018 for a review; Zhao, Wang, & Kim, 2004). In order to remove the effect of ascending veins from the cortical depth fMRI profiles, we employed leakage weights proposed by Markuerkiaga, et al. (2016), and employed a spatial deconvolution approach described in detail in Marquardt et al. (2018). In brief, for each cortical depth level, we subtracted the estimated contribution of all deeper depth levels to obtain an estimate of the ‘true’ local signal change at that depth level.

### Visual field projection

While it is instructive to examine the spatial extent of activation on the inflated cortical surface, the exact relationship between the visual stimulus and the surface activation map is difficult to interpret: Cortical magnification and differences in receptive field size across the cortex complicate the mapping from visual space to the cortical surface. Therefore, we projected the activation maps into the visual field, based on population receptive field estimates. The resulting visual field projections reveal the spatial pattern of activation with respect to the stimulus-space. Population receptive field mapping (Dumoulin & Wandell, 2008) provides three parameters per vertex: x-position, y-position, and size of the Gaussian population receptive field model. For each vertex contained in the ROI, the 2D Gaussian population receptive field model was multiplied with the percent signal change for that vertex. The resulting scaled 2D Gaussians were summed over vertices. The result (a 2D array) was normalised by the population receptive field density at each visual field location (i.e. divided by the sum of 2D Gaussian over vertices). More formally, let ***M****_i,j,k_* be a 3D tensor containing the population receptive field model for visual field positions *i, j* for vertices *k*. The population receptive field model at each visual field location is a 2D Gaussian function:

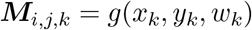

where *x_k_, y_k_, w_k_* are the x-position, y-position, and width (standard deviation) of the 2D Gaussian for vertex *k*, respectively. Further, let ***p****_k_* be a vector with percent signal change values for *n* vertices contained in the ROI. The visual field projection (***V****_i,j_*) of percent signal change values (***p****_k_*) was calculated as:

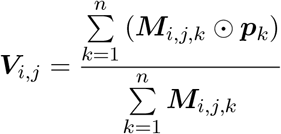

where the multiplication and division operations are element-wise. The visual field projection ***V****_i,j_* was calculated separately for each ROI and cortical depth level, but together for all subjects (by concatenating all subjects’ population receptive field models, ***M****_i,j,k_*, and percent signal change vectors, ***p****_k_*). In this way, all subjects’ activation maps can be projected into a single visual space; this is essentially a simple form of ‘hyperalignment’. (The procedure is similar to that employed by Kok et al. (2016), with the difference that we did not apply any smoothing to the visual field projection.)

### Hypothesis testing

Differences in stimulus-induced activation were investigated by means of a linear mixed effects model. First, we assessed whether the stimuli differentially activated brain areas V1, V2, and V3. (In other words, did activation differ between ROIs as a function of condition?) Second, we tested whether the activation profiles across cortical depth differed between brain areas. Both tests were implemented by means of a mixed effects model including the fixed factors ROI, stimulus condition, and cortical depth, and a random slope for subjects. The autocorrelation structure of cortical depth (within subjects) was modelled as continuous autoregressive of order one. For the first test, a model with all possible two-way interactions was compared with a null model, from which the stimulus condition by ROI interaction had been omitted (because this interaction reflects a differential effect of stimulus condition on brain areas). The second test compared a model with all possible two-way interactions with a null model without the cortical depth by ROI interaction (reflecting differences in cortical depth profiles between areas). The mixed effects models were fitted based on the percent signal change estimate of the sustained and transient predictors (for the stimulus centre and edge, respectively) obtained from the GLM. Comparisons of the respective pairs of models were conducted with a likelihood ratio tests. Models were fitted and compared using R and the nlme package (Pinheiro, Bates, DebRoy, Sarkar, & R Core Team, 2017; R Core Team, 2017).

## Results

In accordance with a previous report using a similar stimulus (Akin et al., 2014), but contrary to what could be the generally expected positive response to a luminance increase, we observed widespread negative signal change in the retinotopic representation of our stimuli in early visual cortex of the right hemisphere. This is illustrated here for the experimental condition inducing the illusory motion percept (‘Pac-Man dynamic’, Figure 2, see also Supplementary Figure S2). A band of positive activation was observed at the cortical representation of the stimulus edge (Figure 2 E, F). The pattern of negative responses to the surface interior and positive activation at the stimulus edge was similar across stimulus conditions (Figure 5; and Supplementary Figure S5).

In the cortical representation of the surface (Figure 2C), we found increased activity due to the illusory percept of motion in the experimental condition, compared to the control conditions where this percept was absent. In particular, the experimental and control conditions showed differential activity with a magnitude that differed among brain areas V1, V2, and V3, as confirmed by a significant ROI (V1, V2, V3) by condition (Pac-Man dynamic, Pac-Man static, control dynamic) interaction (likelihood ratio (df): 39.6 (4), *p* < 0.0001). Moreover, cortical depth profiles of the activity increase were significantly different between brain areas (likelihood ratio (df) of model comparison with/without cortical depth by ROI interaction: 30.2 (2), *p* < 0.0001). Figure 3 shows the cortical depth profile of the signal gain corresponding to the induced motion effect for the cortical representation of the stimulus centre (using the difference between Pac-Mac dynamic and control dynamic). The peak of the apparent motion effect was located at ∼25% in V1, ∼50% in V2, and ∼40% in V3, relative to the pial surface (where 100% cortical depth corresponds to the white/grey matter boundary).

**Figure 3.**
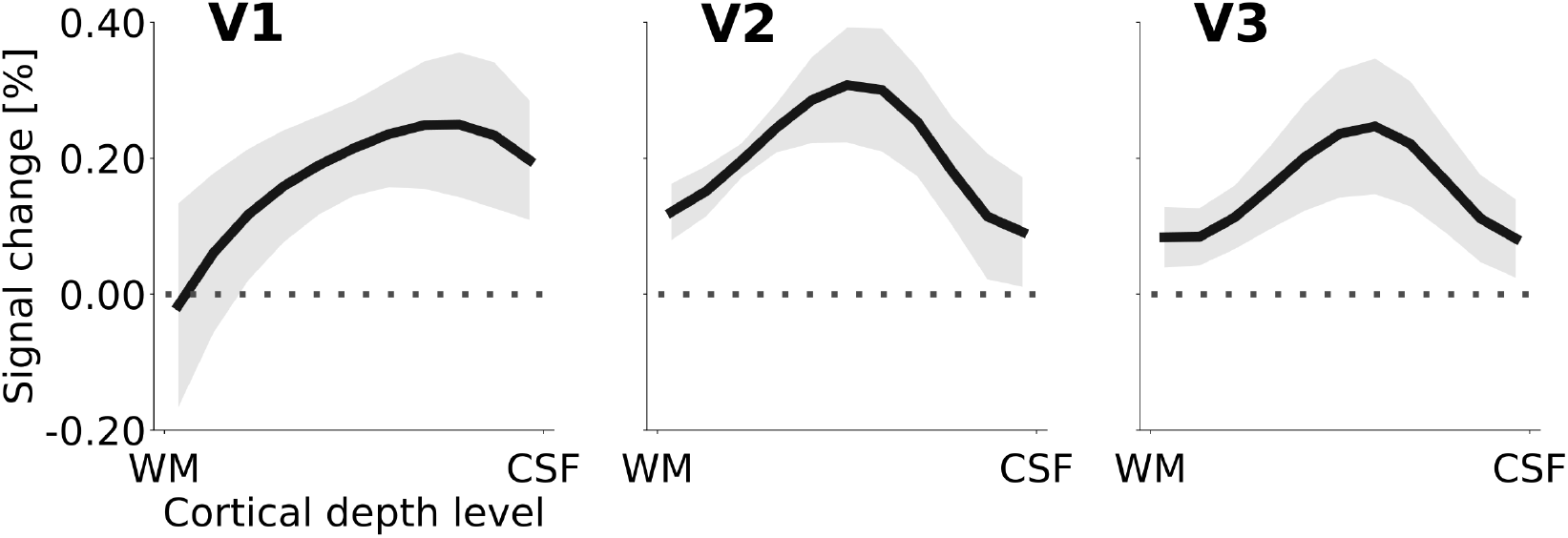
Cortical depth profiles of the apparent motion effect for the cortical representation of the stimulus centre (see Figure 2 B & E). The apparent motion effect was defined as the relative signal change associated with the condition contrast ‘Pac-Man dynamic’ (Figure 1 A) minus ‘control dynamic’ (Figure 1 C). Shading represents the standard error of the mean (across subjects). See Supplementary Figure S3 for the same results for all experimental conditions, and Supplementary Figure S4 for the cortical depth profile of the apparent motion effect at the representation of the stimulus edge.

For the cortical representation of the stimulus edge, the stimulus conditions also caused differential activation among visual areas (likelihood ratio (df) of model comparison with/without ROI by condition interaction: 22.8 (4), *p* < 0.0001). However, there was no evidence for differences between stimulus conditions in the cortical depth profiles at the stimulus edge (likelihood ratio (df) of model comparison with/without cortical depth by condition interaction: 1.6 (2), *p* = 0.46); see Supplementary Figure S4 for cortical depth profiles of apparent motion effect at stimulus edge). This is likely due to the strong feedforward drive due to local contrast at the figure’s edge, which may engage neurons about equally across cortical depth.

### Temporal response pattern

In areas V1, V2, and V3, the central region of interest for all conditions exhibited a sustained negative response, whereas the edge region responded with a transient positive signal change at stimulus onset and offset (Figure 4). Separately for the sustained and transient responses, we determined response onset time as the first time point at which the signal was significantly different from zero (one-sample t-test, *p* < 0.05, Bonferroni corrected). Interestingly, this revealed that the onset of the transient response at the cortical representation of the stimulus edge preceded the onset of the sustained response in the surface representation by one MRI acquisition time point (i.e. ∼2 s; Figure 4). The pattern of positive transient and negative sustained responses at the stimulus edge and centre, respectively, was consistent across areas and conditions (Supplementary Figure S5). An additional control experiment was performed to investigate whether the temporal dynamics of the responses were similar for a longer stimulus duration (Supplementary Figure S6). The results indicate that this was indeed the case, and that the negative response to the centre of the PacMan surface was sustained over long stimulus durations (25 s, compared to ∼10 s in the main experiment).

**Figure 4.**
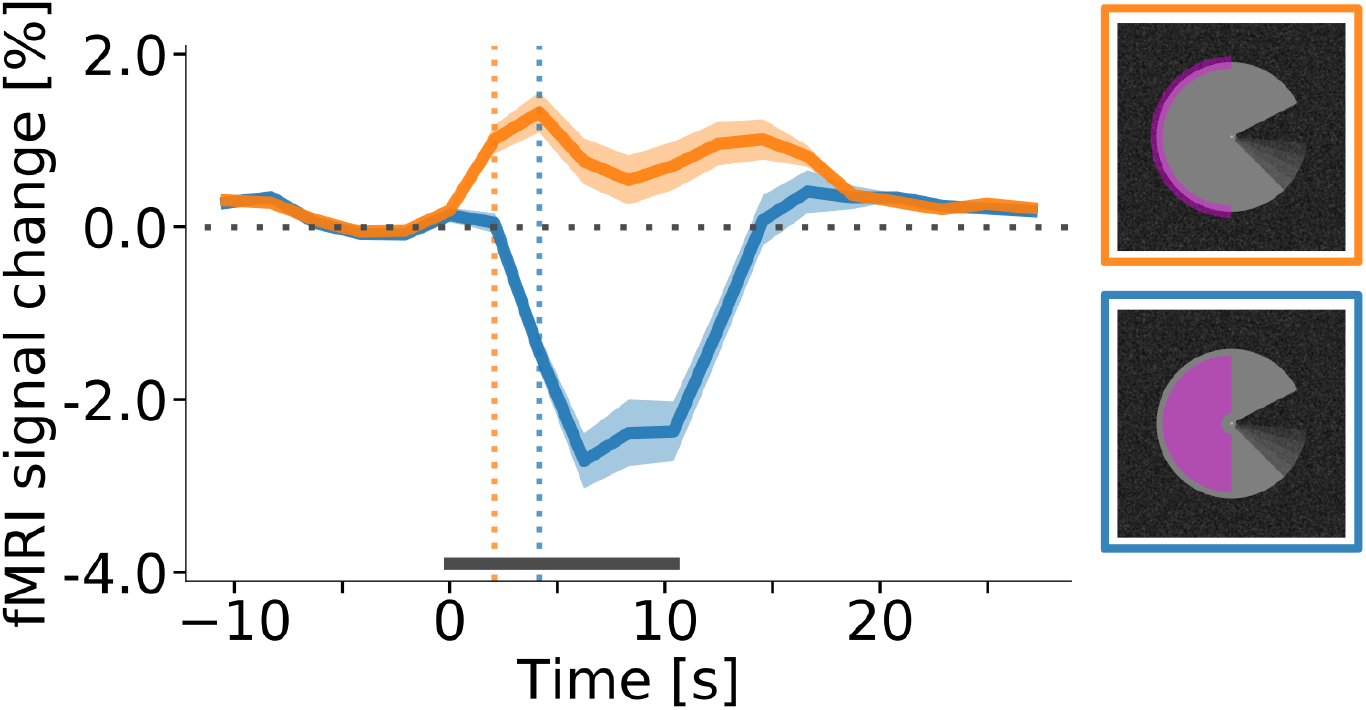
Response onset times in V1. (A) Event-related fMRI timecourses for regions of interest corresponding to the stimulus centre (blue line) and the edge of the stimulus (orange line). The dotted vertical lines indicate the response onset, defined as the first time point at which the signal was significantly different from zero (one-sample t-test, *p* < 0.05, Bonferroni corrected). The positive response at the stimulus edge precedes the negative response at the stimulus centre by one volume (i.e. by about 2 s), suggesting that the negative response is not caused by the onset of the stimulus, but by its prolonged presentation. The response is shown for area V1 of the right hemisphere, averaged (mean) over subjects, stimulus conditions, and cortical depth levels. The horizontal grey bar marks the duration of the stimulus block. Error shading represents the standard error of the mean (across subjects). (See Supplementary Figure S5 for same results separately for all areas and conditions.)

### Spatial response pattern

The spatial distribution of positive and negative signal change is directly visible in the visual field projections (Figure 5). As expected for a moving stimulus, the dynamic parts of the stimulus (i.e. the rotating ‘mouth’ of the Pac-Man, and the rotating wedge of the dynamic control stimulus) caused a positive signal change in their cortical representations in V1, V2, and V3 (Figure 5 A, C, D, F, G, I). All three stimuli caused a negative signal change in the surface’s representation in the right hemisphere in V1, V2 and V3 (Figure 5 A–I). The band of positive signal change seen on the inflated brain (Figure 2 E) is also apparent in the visual field projections (particularly in Figure 5 D, E, F). Especially for the static Pac-Man stimulus, the shape of the stimulus is visible in the visual field projections (Figure 5 B & E), evidence for a high accuracy of the visual field projections across the subjects. The spatial extent of the negative signal change was similar across conditions, but differed across regions; from V1 over V2 to V3, the visual field projections are more blurred, likely due to the increasing neuronal receptive field size in higher-order areas (Gattass, Gross, & Sandell, 1981).

**Figure 5.**
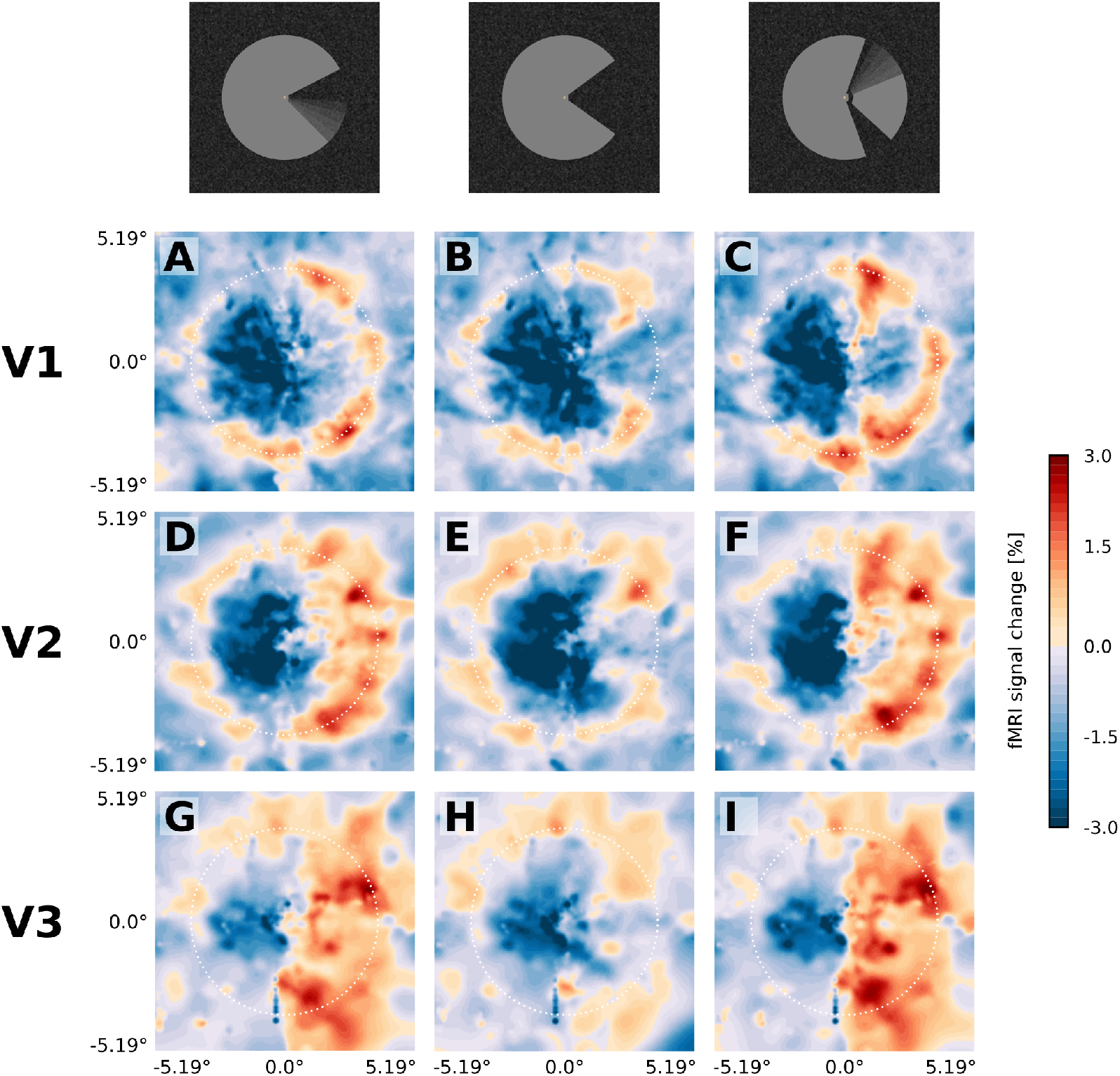
Projection of GLM parameters into visual space. The parameter estimates for the three stimulus conditions (Pac-Man dynamic (A, D, G), Pac-Man static (B, E, H), and control dynamic (C, F, I)) were projected into a model of the visual space based on their retinotopic location, and the size of their respective population receptive fields. The dashed white circles correspond to an eccentricity of 3.75°, i.e. the radius of the Pac-Man stimulus. In all three stimulus conditions, there is a negative response to the left half of the stimulus. Visual field projections are averaged over depth levels (mean).

### Background dependence of the negative response

A control experiment was conducted to investigate the effect of the background and of the stimulus shape on the processing of a surface stimulus. The results revealed that the directionality and temporal course of the response is heavily affected by the type of background, but not by the shape of the stimulus. A negative surface response was only observed when the stimuli were presented on a texture background, irrespective of the stimulus shape (Figure 6 B & D). When presented on a homogenous background, as luminance stimuli are usually presented, the interior of the surface and its edges evoke a positive response (Figure 6 A & C).

The temporal dynamics of the response in the texture background condition (Figure 7, green and blue lines) closely resembled the results from the main experiment (Figure 4); showing a transient positive response at the edges and a sustained, delayed, negative response at the surface interior. In contrast, the response to both the interior of the surface and to its edges was positive and sustained in case of a uniform background (Figure 7, red and orange lines).

**Figure 6.**
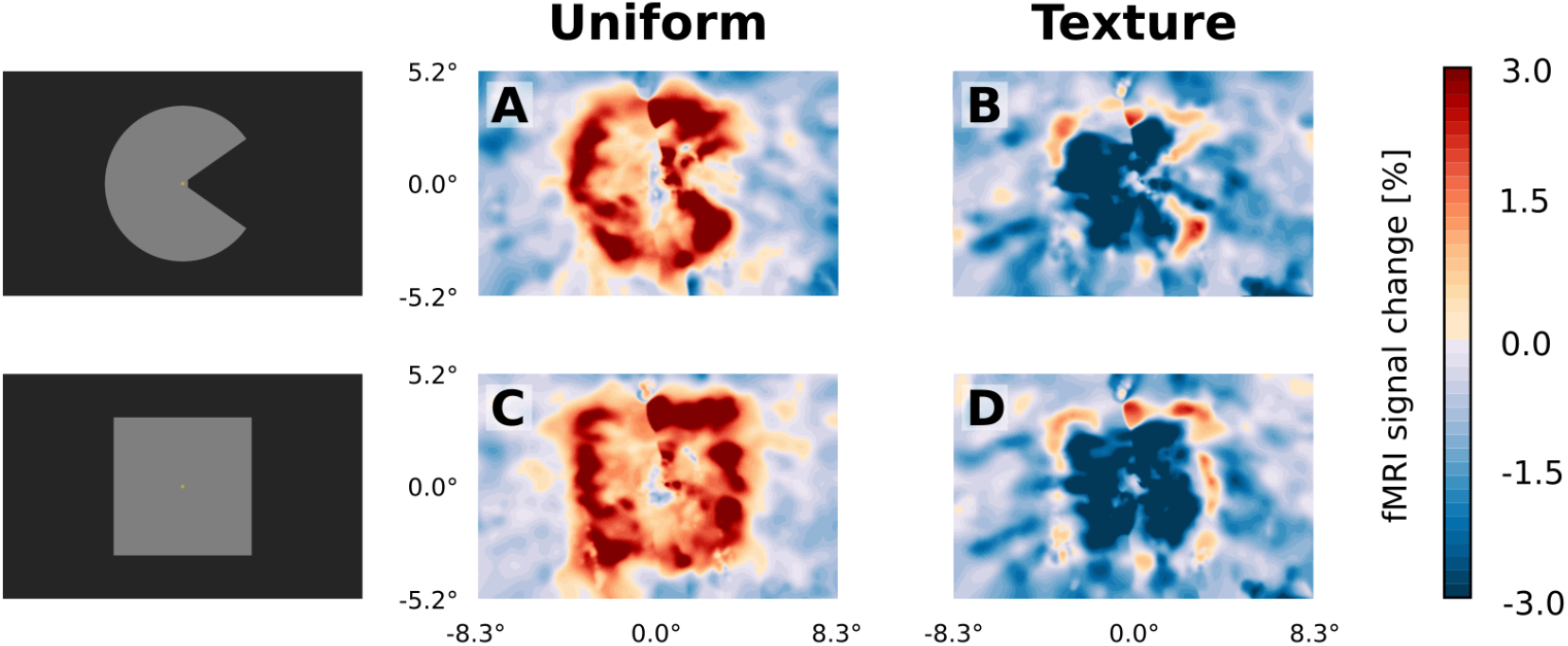
Visual field projections of GLM parameter estimates from control experiment with texture background and uniform background, for V1. A ‘Pac-Man’ figure and a square were presented either on a uniform background (A & C) or on a random texture background (B & D). When presented on a uniform background, the stimuli caused a positive response, especially at the retinotopic representation of the edges (A & C). In stark contrast, the response to the interior of the stimuli was negative when presented on a random texture background (B & D). At the edges of the stimuli, a small band of positive activity can still be observed (B & D).

**Figure 7.**
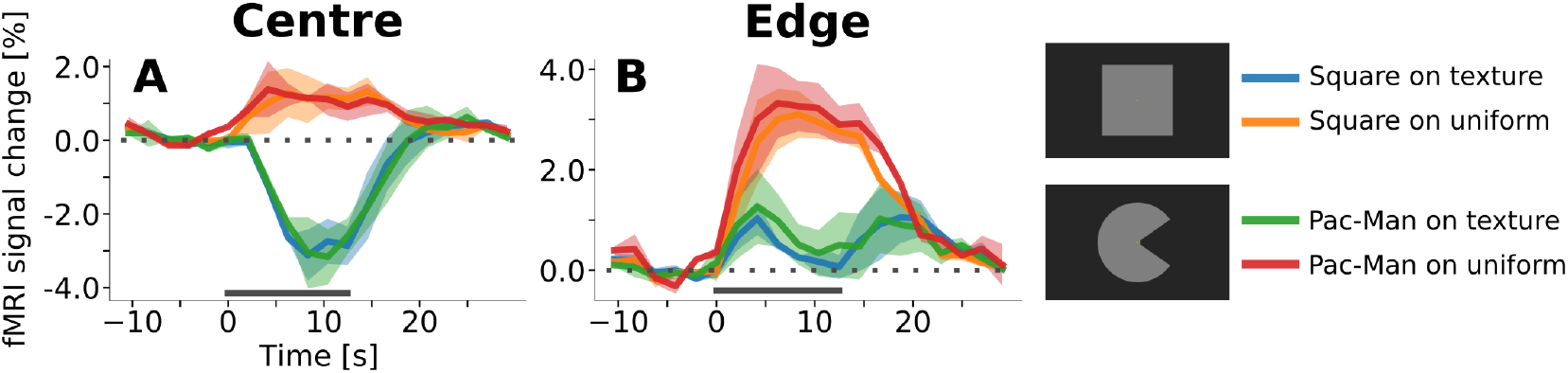
Event-related time courses from control experiment with texture background and uniform background, separately for regions of interest corresponding to the retinotopic representation of the centre of the stimulus (A) and to its edges (B). Irrespective of the shape of the stimulus (square or ‘Pac-Man’), there is a positive response to the centre of the stimulus when the background is uniform (A, red & orange lines), and a negative response when the stimuli are presented on a random texture pattern (A, green & blue lines). Interestingly, the positive response has a shorter latency than the negative response. The response to the edges of the stimuli is positive under all conditions (B). However, the response amplitude is much stronger when the stimuli are presented on a uniform background. Moreover, the temporal dynamics changes as a function of the background; the response is sustained when the background is uniform (B, orange & red lines), but transient for the texture background (B, green and blue lines). The horizontal grey bar marks the duration of the stimulus.

Interestingly, these results imply that the temporal shape of the edge response changed as a function of the background condition; in other words, whether the edge response is sustained or transient depends on whether the stimuli are presented on a texture pattern or on a uniform background.

## Discussion

We have studied neural correlates of perceived surface motion induced in a locally static grey surface on a dark, textured background (Figure 1). The motion percept was caused by local edge movement in the contralateral hemifield and spread over the entire surface in the ipsilateral hemifield. We report three main findings: First, the induced percept of surface motion was associated with an fMRI signal increase in the representation of the surface in areas V1, V2 and V3 (Figure 3). As the enhanced signal was measured far away from the location where the perceived motion was induced, this signal likely derives from feedback. In addition, the differences in the cortical depth distribution of motion-percept related signal gain among visual areas also supported a feedback origin. Second, we found that the response to the edge preceded the response to the surface by approximately 2 s (Figure 4). Third, we observed a negative BOLD signal in the figure representation (Figure 5), which depended on the presence of a textured background and was eliminated when the background texture was removed (Figure 6). Hence, the signal gain due to the motion percept represented an increase in signal from a negative BOLD signal in the control condition to a less negative BOLD signal in the illusory movement-condition.

### Top-down feedback

The main and control stimuli were ‘physically’ identical in the left visual field, while the global perceptual quality of the stimulus depended on the right half of the stimuli (Figure 1; videos of the stimuli are available online: https://doi.org/10.5281/zenodo.2583017). This stimulus design offers three advantages: First, the surface itself was homogenously grey and did not contain local moving elements, thereby avoiding the interpretative question whether enhanced fMRI activity during surface perception reflects enhanced processing of local elements or an integrated surface motion percept. Second, any changes in activity correlating with a perceptual change from static to moving in the left hemifield were induced by the right hemifield. Anatomical investigations have shown that transcallosal, interhemispheric connections are restricted to the proximity of the vertical meridian in primate early visual cortex (Clarke & Miklossy, 1990; Essen & Zeki, 1978; Glickstein & Whitteridge, 1976; Wong-Riley, 1974). This, combined with the fact that the surface motion percept in the left hemifield was induced in the absence of physical changes to the left-hemifield stimulus, renders top-down feedback from higher areas, rather than within-area horizontal interactions, the most plausible source of the motion percept and associated depth distributions of activity. Third, the cortical region that retinotopically represents the physically constant left side of the stimulus and the one which induces the motion percept (i.e. the ‘mouth’ of the Pac-Man) were far apart. Thus, it is very unlikely that imprecisions in the retinotopic maps could confound our results. By the same token, the size of our stimulus enabled us to separate responses to the surface from responses to the contours.

The cortical depth profiles of the enhanced response due to the illusory motion effect in V1, V2, and V3 suggests that top-down signals may have re-entered at superficial layers in V1, where most of the signal gain due to motion perception is concentrated (Figure 3). Re-entrant connections via superficial V1 have been reported in neurophysiological (McManus, Li, & Gilbert, 2011), anatomical (Martinez-Conde et al., 1999), and high-field fMRI studies (Muckli et al., 2015). This re-entrant information may have propagated to V2 and V3 via feedforward pathways, in line with anatomical evidence that the strongest forward projections from V1 to V2 originate in superficial V1 layers 3B and 4B, and arrive across the full extent of layer 4 in V2 (Douglas & Martin, 2004; Felleman & Van Essen, 1991). Furthermore, forward projections originating in superficial V1 layers and superficial V2 layers also target layer 4 in V3 (Rockland & Pandya, 1979; Van Essen, Newsome, Maunsell, & Bixby, 1986). This pattern of forward projections may explain the activity peak at intermediate depths of areas V2 and V3 (Figure 8A). Therefore, although our data do not permit a direct test of the directionality and precise temporal dynamics of information flow, re-entrant feedback at the level of V1 is a plausible interpretation of the present results.

**Figure 8.**
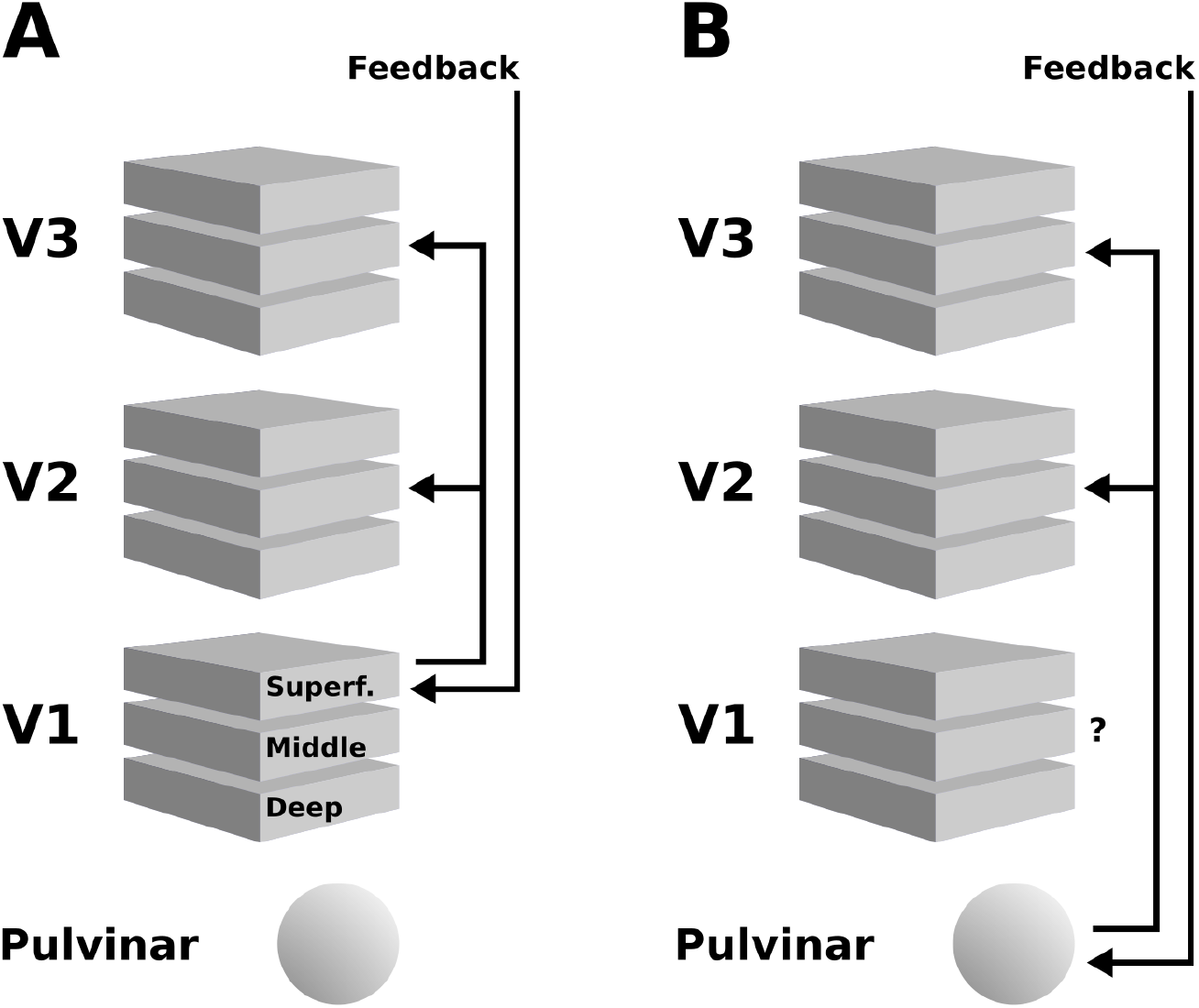
Schematic illustration of two possible interpretations of the present results. (A) Higher cortical areas may integrate the global motion percept across hemispheres, and send feedback projections to superficial layers of V1. Subsequently, this re-entrant feedback would be sent to V2 and V3 via feedforward connections. (B) Alternatively, the pulvinar may act as a ‘higher-order relay’, and send feedback from higher cortical areas to V2 and V3. See discussion section for details.

An additional contribution to the depth-pattern of activity observed in extrastriate areas may have originated from the pulvinar, and possibly other subcortical structures (Standage & Benevento, 1983; Trojanowski & Jacobson, 1977). The middle layers of extrastriate cortex are the target of projections from the pulvinar (Benevento & Rezak, 1976; Figure 8B Benevento, Rezak, & Bos, 1975; Ogren & Hendrickson, 1977; Rezak & Benevento, 1979), a structure that is sometimes referred to as a ‘higher-order relay’ because of its role in cortico-cortical interaction (Sherman & Guillery, 2002). The pulvinar has been shown to regulate corticocortical communication in the visual system based on attentional demands (Saalmann, Pinsk, Wang, Li, & Kastner, 2012). Experiments in humans (Villeneuve, Kupers, Gjedde, Ptito, & Casanova, 2005; Villeneuve, Thompson, Hess, & Casanova, 2012) and cats (Merabet, Desautels, Minville, & Casanova, 1998) have demonstrated a role of the pulvinar in higher-order motion processing (i.e. coherent motion of entire objects, as opposed to local motion). In line with this, Shimono et al. have found evidence for an involvement of the pulvinar in the interhemispheric integration of motion information (2012). In summary, both cortical and subcortical sources of re-entrant feedback in lower-level visual areas may have contributed to the observed depth-resolved responses (see Figure 8).

The positive BOLD contribution associated with the illusory percept of surface motion is in line with other fMRI studies for a range of surface illusions (Hsieh & Tse, 2010; Kok & de Lange, 2014; Mendola et al., 1999; Pereverzeva & Murray, 2008; Sasaki & Watanabe, 2004). Compared to (Kok et al., 2016), who reported a fMRI response enhancement limited to the deepest cortical layers during the percept of an illusory Kansiza triangle, the signal gain we found was focused on superficial to middle layer compartments. Our results resemble somewhat more the superficial activity reported in (Muckli et al., 2015) in response to the completion of occluded visual scenes. These differences in activity depth profiles could reflect fundamental differences in feedback mechanisms engaged in the stimulus paradigms in the different studies, which is a possibility that should be investigated further. Irrespective of the differences in observed activity profiles over depth, they all support re-entrant feedback signals, which is in line with mounting evidence that, even for the simplest displays, feedback from the highest level of the visual system plays a role (Lamme & Roelfsema, 2000; McManus et al., 2011; Roelfsema, Lamme, Spekreijse, & Bosch, 2002; Schnabel et al., 2018).

### Edge responses preceding surface responses

Psychophysical experiments (Paradiso & Nakayama, 1991) and neurophysiological experiments (Huang & Paradiso, 2008) have suggested that surface brightness may fill in from the edge over a time period of ∼100 ms, depending on the size of the surface. This interpretation of the reported data is in line with computational models that propose a primary analysis of the visual scene to delineate contours, followed by a secondary analysis that is initiated by and interacts with these contours to reconstruct the visible aspect of the surfaces (Grossberg & Hong, 2006; Pessoa, Mingolla, & Neumann, 1995). Although these models have proposed diffusion-like processes in retinotopic visual areas as a neural correlate for surface perception, feedback processes related to surface processing also display a delayed modulation of activity in early visual cortex of >100 ms (Lamme et al., 1999; Self et al., 2013). In addition, low-level aspects of the stimulus, such as the enhanced contrast at the edge and the absence of contrast inside the grey figure, can induce faster response latencies in early visual cortex at the edge representation compared to inside the homogeneous figure (Albrecht, Geisler, Frazor, & Crane, 2002). Conceptually, an initial analysis of edges can also be seen as generating predictions for the presence of surfaces and their features, in line with the predictive coding hypothesis (Rao & Ballard, 1999). Hence, the earlier response to the edge compared to the surface is generally in line with a range of existing concepts and data about surface perception, but the question is whether and how this small temporal difference in neuronal responses translates into a ∼2 s difference in BOLD response onset (see Figure 4). It is possible that the apparent delay in the onset of the BOLD response to the surface may be the result of competing positive and negative BOLD effects (Uludağ & Blinder, 2018). In the surface cortical representation, positive (due to luminance increase) and negative (due to lateral inhibition) BOLD responses may occur equally quickly and strongly, and hence may balance each other at the beginning of the stimulation. As time passes, the negative response may appear due to a more sustained negative response paired with a more transient or adaptive positive response. Thus, even though both the positive and negative BOLD responses may have similar latencies as the edge response, the sum of both centre responses may initially cancel out and lead to a larger apparent latency of the negative response emerging later on.

### Negative BOLD response

In contrast to an expected increase of the BOLD signal in response to a local luminance increase, but in line with a previous study (Akin et al., 2014), the surfaces yielded strongly negative BOLD responses in V1, V2 and V3, irrespective of whether they were perceived as static or moving (Figure 5). The negative response was located at the cortical retinotopic representation of the interior of the surface and was sustained throughout the presentation period (Figure 4, and Supplementary Figures S5 & S6). Note that the effect of the background on the fMRI signal related to the surface area is very strong. A change in the background from texture to homogeneous dark background resulted in a 4% signal change (from −3% to +1% BOLD). It is quite remarkable that a subtle change in the background leads to such a strong decrease in BOLD signal and presumably reduction in metabolism and excitatory neuronal activity. In comparison, Kok et al. observed a response amplitude of approximately 0.7% to 1.4% at the retinotopic representation of a centrally presented contrast-reversing checkerboard (using a similar MRI pulse sequence and the same spatial resolution as in the present study, Kok et al., 2016, see their Figure S2 B). Previous studies have linked negative BOLD with a inhibitory competition between large, juxtaposed stimulated and unstimulated regions (Shmuel, Augath, Oeltermann, & Logothetis, 2006; Shmuel et al., 2002).

In our experiment, we speculate that at the boundary, as well as within the figure and the textured background region, there is a competition between inhibitory and excitatory processes that results in the observed patterns of positive and negative BOLD responses. Although at present we do not understand the underlying mechanisms, the fact that extreme changes in the patterns of negative and positive BOLD signals depend on the presence of a very subtle texture in the background, suggests a determining influence of feedback signals. The fact that the response to the edge of the figure rises faster than the response to the surface (Figures 4 and 7) may reflect differences in the speed at which the hypothesized competition is settled at the figure’s edges and within its interior.

### Spatial deconvolution

A complicating factor in the analysis of the layered distribution of fMRI signal is related to the anatomy of ascending draining veins, which leads to a strong bias for the BOLD signal to be stronger in superficial cortical layers, even if the neuronal activity is stronger in deeper layers (Koopmans et al., 2011; Markuerkiaga et al., 2016; see Uludağ & Blinder, 2018 for a review). To use the BOLD signal as a realistic estimate of underlying neural activity in high-resolution data, it is therefore crucial to take this effect into account (Markuerkiaga et al., 2016). We have previously employed a spatial deconvolution to remove signal spread due to ascending veins (Marquardt et al., 2018). The exact parameters of the spatial deconvolution are difficult to determine, and our parameter choices may not be exact. Nevertheless, simulations have shown that the spatial deconvolution is relatively robust against deviations in its model parameters (see Marquardt et al., 2018, Figure 8, and Supplementary Figures S4 & S5 therein). Although the exact shape of the resulting cortical depth profiles is contingent on the model parameters of the spatial deconvolution, the results do not differ qualitatively in case of different model parameters within physiologically plausible ranges (Marquardt et al., 2018). Thus, we stress the importance of data analysis, in general, and spatial deconvolution, in particular, for high-resolution fMRI to obtaining accurate representation of neuronal activity across cortical depths.

### Summary

Our study provides the first evidence that a motion percept in a surface region of a stimulus far removed from the local information inducing the motion percept produces a small increase in activity in the retinotopic representation of the figure. At the same time, our study reports a negative BOLD signal in the figure representation of an unexpected magnitude, and in contrast to standard expectation, following a luminance increase. This shows that subtle low-level aspects of the stimulus can have pronounced effects not only on the magnitude but even on the sign of the BOLD signal. It is an open question whether the neural mechanisms behind the negative response have a functional role in surface perception. In spite of the negative BOLD response, the perceptual assignment of a surface feature to a visual field region (where that feature was physically absent) yielded a signal enhancement, in line with other studies. While different surface features or displays may result in distinct depth resolved patterns of fMRI activity, possibly suggesting various sources of feedback, the consistent finding of signal enhancements during induced or illusory surface perception also suggests common aspects to the mechanisms of surface perception independent of the displays or features.

## Acknowledgments

This work was financially supported by funding from IBS (#IBS-R015-D1) to KU and the Netherlands Organization for Scientific Research (NWO; Research Talent 406-14-085) to KU and IM.

## Author Contributions

I.M., M.S., O.F.G., and K.U. conceptualized research; I.M., M.S., and D.I. conducted the investigation; I.M., M.S., O.F.G., and K.U. designed methodology; I.M., M.S., and O.F.G. contributed software; I.M. curated the data; I.M. performed formal analysis; K.U. and I.M. acquired funding, K.U. and P.d.W. supervised the project; I.M., P.d.W., and K.U. prepared, reviewed, and edited the paper.

## References

Ahrens, J., Geveci, B., & Law, C. (2005). ParaView: An End-User Tool for Large-Data Visualization. In C. D. Hansen & C. R. Johnson, The Visualization Handbook (pp. 717–731). Burlington: Elsevier Butterworth–Heinemann.

Akin, B., Ozdem, C., Eroglu, S., Keskin, D. T., Fang, F., Doerschner, K., … Boyaci, H. (2014). Attention modulates neuronal correlates of interhemispheric integration and global motion perception. Journal of Vision, 14(12). https://doi.org/10.1167/14.12.30

Albrecht, D. G., Geisler, W. S., Frazor, R. A., & Crane, A. M. (2002). Visual cortex neurons of monkeys and cats: temporal dynamics of the contrast response function. Journal of Neurophysiology, 88(2), 888–913.

Albrecht, D. G., & Hamilton, D. B. (1982). Striate cortex of monkey and cat: contrast response function. Journal of Neurophysiology, 48(1), 217–237.

Anderson, J. C., & Martin, K. A. C. (2009). The synaptic connections between cortical areas V1 and V2 in macaque monkey. The Journal of Neuroscience: The Official Journal of the Society for Neuroscience, 29(36), 11283–11293. https://doi.org/10.1523/JNEUROSCI.5757-08.2009

Andersson, J. L. R., Skare, S., & Ashburner, J. (2003). How to correct susceptibility distortions in spin-echo echo-planar images: application to diffusion tensor imaging. NeuroImage, 20(2), 870–888. https://doi.org/10.1016/S1053-8119(03)00336-7

Ayachit, U. (2015). The ParaView guide: updated for ParaView version 4.3 (Full color version; L. Avila, Ed.). Los Alamos: Kitware.

Bazin, P.-L., Cuzzocreo, J. L., Yassa, M. A., Gandler, W., McAuliffe, M. J., Bassett, S. S., & Pham, D. L. (2007). Volumetric neuroimage analysis extensions for the MIPAV software package. Journal of Neuroscience Methods, 165(1), 111–121. https://doi.org/10.1016/j.jneumeth.2007.05.024

Behnel, S., Bradshaw, R., Citro, C., Dalcin, L., Seljebotn, D. S., & Smith, K. (2011). Cython: The Best of Both Worlds. Computing in Science & Engineering, 13(2), 31–39. https://doi.org/10.1109/MCSE.2010.118

Benevento, L. A., & Rezak, M. (1976). The cortical projections of the inferior pulvinar and adjacent lateral pulvinar in the rhesus monkey (Macaca mulatta): an autoradiographic study. Brain Research, 108(1), 1–24.

Benevento, L. A., Rezak, M., & Bos, J. (1975). Extrageniculate projections to layers VI and I of striate cortex (area 17) in the rhesus monkey (Macaca mulatta). Brain Research, 96(1), 51–55.

Clarke, S., & Miklossy, J. (1990). Occipital cortex in man: organization of callosal connections, related myelo- and cytoarchitecture, and putative boundaries of functional visual areas. The Journal of Comparative Neurology, 298(2), 188–214. https://doi.org/10.1002/cne.902980205

Cornelissen, F. W., Wade, A. R., Vladusich, T., Dougherty, R. F., & Wandell, B. A. (2006). No functional magnetic resonance imaging evidence for brightness and color filling-in in early human visual cortex. The Journal of Neuroscience: The Official Journal of the Society for Neuroscience, 26(14), 3634–3641. https://doi.org/10.1523/JNEUROSCI.4382-05.2006

De Weerd, P., Gattass, R., Desimone, R., & Ungerleider, L. G. (1995). Responses of cells in monkey visual cortex during perceptual filling-in of an artificial scotoma. Nature, 377(6551), 731–734. https://doi.org/10.1038/377731a0

Dennett, D. C. (1991). Consciousness explained (1st ed). Boston: Little, Brown and Co.

Douglas, R. J., & Martin, K. A. C. (2004). Neuronal circuits of the neocortex. Annual Review of Neuroscience, 27, 419–451. https://doi.org/10.1146/annurev.neuro.27.070203.144152

Dumoulin, S. O., & Wandell, B. A. (2008). Population receptive field estimates in human visual cortex. NeuroImage, 39(2), 647–660. https://doi.org/10.1016/j.neuroimage.2007.09.034

Essen, D. C., & Zeki, S. M. (1978). The topographic organization of rhesus monkey prestriate cortex. The Journal of Physiology, 277, 193–226.

Feinberg, D. A., Moeller, S., Smith, S. M., Auerbach, E., Ramanna, S., Gunther, M., … Yacoub, E. (2010). Multiplexed echo planar imaging for sub-second whole brain FMRI and fast diffusion imaging. PloS One, 5(12), e15710. https://doi.org/10.1371/journal.pone.0015710

Felleman, D. J., & Van Essen, D. C. (1991). Distributed hierarchical processing in the primate cerebral cortex. Cerebral Cortex (New York, N.Y.: 1991), 1(1), 1–47.

Friedman, H. S., Zhou, H., & von der Heydt, R. (2003). The coding of uniform colour figures in monkey visual cortex. The Journal of Physiology, 548(Pt 2), 593–613. https://doi.org/10.1113/jphysiol.2002.033555

Friston, K. J., Williams, S., Howard, R., Frackowiak, R. S., & Turner, R. (1996). Movement-related effects in fMRI time-series. Magnetic Resonance in Medicine, 35(3), 346–355.

Gattass, R., Gross, C. G., & Sandell, J. H. (1981). Visual topography of V2 in the macaque. The Journal of Comparative Neurology, 201(4), 519–539. https://doi.org/10.1002/cne.902010405

Glickstein, M., & Whitteridge, D. (1976). Degeneration of layer III pyramidal cells in area 18 following destruction of callosal input. Brain Research, 104(1), 148–151.

Gregory, R. L. (1972). Cognitive contours. Nature, 238(5358), 51–52.

Greve, D. N., & Fischl, B. (2009). Accurate and robust brain image alignment using boundary-based registration. NeuroImage, 48(1), 63–72. https://doi.org/10.1016/j.neuroimage.2009.06.060

Grosof, D. H., Shapley, R. M., & Hawken, M. J. (1993). Macaque V1 neurons can signal ‘illusory’ contours. Nature, 365(6446), 550–552. https://doi.org/10.1038/365550a0

Grossberg, S. (1987a). Cortical dynamics of three-dimensional form, color, and brightness perception: I. Monocular theory. Perception & Psychophysics, 41(2), 87–116.

Grossberg, S. (1987b). Cortical dynamics of three-dimensional form, color, and brightness perception: II. Binocular theory. Perception & Psychophysics, 41(2), 117–158.

Grossberg, S., & Hong, S. (2006). A neural model of surface perception: lightness, anchoring, and filling-in. Spatial Vision, 19(2–4), 263–321.

Guidi, M., Huber, L., Lampe, L., Gauthier, C. J., & Möller, H. E. (2016). Lamina-dependent calibrated BOLD response in human primary motor cortex. NeuroImage, 141, 250–261. https://doi.org/10.1016/j.neuroimage.2016.06.030

Gulban, O. F., Schneider, M., Marquardt, I., Haast, R. A. M., & De Martino, F. (2018). A scalable method to improve gray matter segmentation at ultra high field MRI. PloS One, 13(6), e0198335. https://doi.org/10.1371/journal.pone.0198335

Halchenko, Y. O., & Hanke, M. (2012). Open is Not Enough. Let’s Take the Next Step: An Integrated, Community-Driven Computing Platform for Neuroscience. Frontiers in Neuroinformatics, 6, 22. https://doi.org/10.3389/fninf.2012.00022

Horiguchi, H., Nakadomari, S., Misaki, M., & Wandell, B. A. (2009). Two temporal channels in human V1 identified using fMRI. NeuroImage, 47(1), 273–280. https://doi.org/10.1016/j.neuroimage.2009.03.078

Hsieh, P.-J., & Tse, P. U. (2010). ‘Brain-reading’ of perceived colors reveals a feature mixing mechanism underlying perceptual filling-in in cortical area V1. Human Brain Mapping, 31(9), 1395–1407. https://doi.org/10.1002/hbm.20946

Huang, X., & Paradiso, M. A. (2008). V1 response timing and surface filling-in. Journal of Neurophysiology, 100(1), 539–547. https://doi.org/10.1152/jn.00997.2007

Hubel, D. H., & Wiesel, T. N. (1968). Receptive fields and functional architecture of monkey striate cortex. The Journal of Physiology, 195(1), 215–243. https://doi.org/10.1113/jphysiol.1968.sp008455

Huber, L., Goense, J. B. M., Kennerley, A. J., Trampel, R., Guidi, M., Reimer, E., … Möller, H. E.. (2015). Cortical lamina-dependent blood volume changes in human brain at 7 T. NeuroImage, 107, 23–33. https://doi.org/10.1016/j.neuroimage.2014.11.046

Hunter, J. D. (2007). Matplotlib: A 2D Graphics Environment. Computing in Science & Engineering, 9(3), 90–95. https://doi.org/10.1109/MCSE.2007.55

Jenkinson, M., Bannister, P., Brady, M., & Smith, S. (2002). Improved optimization for the robust and accurate linear registration and motion correction of brain images. NeuroImage, 17(2), 825–841.

Jenkinson, M., & Smith, S. (2001). A global optimisation method for robust affine registration of brain images. Medical Image Analysis, 5(2), 143–156.

Kaczmarzyk, J., Goncalves, M., Halchenko, Y., Mitchell, R., Nielson, D., Jarecka, D., … Rokem, A. (2017, November 18). Kaczmarj/Neurodocker: Version 0.3.2. https://doi.org/10.5281/zenodo.1058998

Keil, M. S., Cristóbal, G., Hansen, T., & Neumann, H. (2005). Recovering real-world images from single-scale boundaries with a novel filling-in architecture. Neural Networks: The Official Journal of the International Neural Network Society, 18(10), 1319–1331. https://doi.org/10.1016/j.neunet.2005.04.003

Kok, P., Bains, L. J., van Mourik, T., Norris, D. G., & de Lange, F. P. (2016). Selective Activation of the Deep Layers of the Human Primary Visual Cortex by Top-Down Feedback. Current Biology, 26(3), 371–376. https://doi.org/10.1016/j.cub.2015.12.038

Kok, P., & de Lange, F. P. (2014). Shape perception simultaneously up- and downregulates neural activity in the primary visual cortex. Current Biology: CB, 24(13), 1531–1535. https://doi.org/10.1016/j.cub.2014.05.042

Komatsu, H., Kinoshita, M., & Murakami, I. (2000). Neural responses in the retinotopic representation of the blind spot in the macaque V1 to stimuli for perceptual filling-in. The Journal of Neuroscience: The Official Journal of the Society for Neuroscience, 20(24), 9310–9319.

Koopmans, P. J., Barth, M., & Norris, D. G. (2010). Layer-specific BOLD activation in human V1. Human Brain Mapping, 31(9), 1297–1304. https://doi.org/10.1002/hbm.20936

Koopmans, P. J., Barth, M., Orzada, S., & Norris, D. G. (2011). Multi-echo fMRI of the cortical laminae in humans at 7T. NeuroImage, 56(3), 1276–1285. https://doi.org/10.1016/j.neuroimage.2011.02.042

Lamme, V. A. (1995). The neurophysiology of figure-ground segregation in primary visual cortex. The Journal of Neuroscience: The Official Journal of the Society for Neuroscience, 15(2), 1605–1615.

Lamme, V. A., Rodriguez-Rodriguez, V., & Spekreijse, H. (1999). Separate processing dynamics for texture elements, boundaries and surfaces in primary visual cortex of the macaque monkey. Cerebral Cortex (New York, N.Y.: 1991), 9(4), 406–413.

Lamme, V. A., & Roelfsema, P. R. (2000). The distinct modes of vision offered by feedforward and recurrent processing. Trends in Neurosciences, 23(11), 571–579.

Lee, T. S., & Nguyen, M. (2001). Dynamics of subjective contour formation in the early visual cortex. Proceedings of the National Academy of Sciences of the United States of America, 98(4), 1907–1911. https://doi.org/10.1073/pnas.031579998

Lu, H. D., & Roe, A. W. (2007). Optical imaging of contrast response in Macaque monkey V1 and V2. Cerebral Cortex (New York, N.Y.: 1991), 17(11), 2675–2695. https://doi.org/10.1093/cercor/bhl177

Markuerkiaga, I., Barth, M., & Norris, D. G. (2016). A cortical vascular model for examining the specificity of the laminar BOLD signal. NeuroImage, 132, 491–498. https://doi.org/10.1016/j.neuroimage.2016.02.073

Marquardt, I., Gulban, O. F., & Schneider, M. (2018, April 18). pyprf - A free & open source python package for population receptive field (pRF) analysis. https://doi.org/10.5281/zenodo.1220207

Marquardt, I., Schneider, M., Gulban, O. F., Ivanov, D., & Uludağ, K. (2018). Cortical depth profiles of luminance contrast responses in human V1 and V2 using 7 T fMRI. Human Brain Mapping. https://doi.org/10.1002/hbm.24042

Marques, J. P., Kober, T., Krueger, G., van der Zwaag, W., Van de Moortele, P.-F., & Gruetter, R. (2010). MP2RAGE, a self bias-field corrected sequence for improved segmentation and T1-mapping at high field. NeuroImage, 49(2), 1271–1281. https://doi.org/10.1016/j.neuroimage.2009.10.002

Martinez-Conde, S., Cudeiro, J., Grieve, K. L., Rodriguez, R., Rivadulla, C., & Acuña, C. (1999). Effects of feedback projections from area 18 layers 2/3 to area 17 layers 2/3 in the cat visual cortex. Journal of Neurophysiology, 82(5), 2667–2675. https://doi.org/10.1152/jn.1999.82.5.2667

McManus, J. N. J., Li, W., & Gilbert, C. D. (2011). Adaptive shape processing in primary visual cortex. Proceedings of the National Academy of Sciences of the United States of America, 108(24), 9739–9746. https://doi.org/10.1073/pnas.1105855108

Mendola, J. D., Dale, A. M., Fischl, B., Liu, A. K., & Tootell, R. B. (1999). The representation of illusory and real contours in human cortical visual areas revealed by functional magnetic resonance imaging. The Journal of Neuroscience: The Official Journal of the Society for Neuroscience, 19(19), 8560–8572.

Meng, M., Remus, D. A., & Tong, F. (2005). Filling-in of visual phantoms in the human brain. Nature Neuroscience, 8(9), 1248–1254. https://doi.org/10.1038/nn1518

Merabet, L., Desautels, A., Minville, K., & Casanova, C. (1998). Motion integration in a thalamic visual nucleus. Nature, 396(6708), 265–268. https://doi.org/10.1038/24382

Millman, K. J., & Aivazis, M. (2011). Python for Scientists and Engineers. Computing in Science & Engineering, 13(2), 9–12. https://doi.org/10.1109/MCSE.2011.36

Moeller, S., Yacoub, E., Olman, C. A., Auerbach, E., Strupp, J., Harel, N., & Uğurbil, K. (2010). Multiband multislice GE-EPI at 7 tesla, with 16-fold acceleration using partial parallel imaging with application to high spatial and temporal whole-brain fMRI. Magnetic Resonance in Medicine, 63(5), 1144–1153. https://doi.org/10.1002/mrm.22361

Muckli, L., De Martino, F., Vizioli, L., Petro, L. S., Smith, F. W., Ugurbil, K., … Yacoub, E. (2015). Contextual Feedback to Superficial Layers of V1. Current Biology : CB, 25(20), 2690–2695. https://doi.org/10.1016/j.cub.2015.08.057

Muckli, L., Kohler, A., Kriegeskorte, N., & Singer, W. (2005). Primary visual cortex activity along the apparent-motion trace reflects illusory perception. PLoS Biology, 3(8), e265. https://doi.org/10.1371/journal.pbio.0030265

Muckli, L., Singer, W., Zanella, F. E., & Goebel, R. (2002). Integration of multiple motion vectors over space: an fMRI study of transparent motion perception. NeuroImage, 16(4), 843–856.

Ogren, M. P., & Hendrickson, A. E. (1977). The distribution of pulvinar terminals in visual areas 17 and 18 of the monkey. Brain Research, 137(2), 343–350.

Oliphant, T. E. (2007). Python for Scientific Computing. Computing in Science & Engineering, 9(3), 10–20. https://doi.org/10.1109/MCSE.2007.58

Olman, C. A., Harel, N., Feinberg, D. A., He, S., Zhang, P., Ugurbil, K., & Yacoub, E. (2012). Layer-specific fMRI reflects different neuronal computations at different depths in human V1. PloS One, 7(3), e32536. https://doi.org/10.1371/journal.pone.0032536

Paradiso, M. A., & Nakayama, K. (1991). Brightness perception and filling-in. Vision Research, 31(7–8), 1221–1236.

Peirce, J. W. (2007). PsychoPy–Psychophysics software in Python. Journal of Neuroscience Methods, 162(1–2), 8–13. https://doi.org/10.1016/j.jneumeth.2006.11.017

Peirce, J. W. (2008). Generating stimuli for neuroscience using PsychoPy. Frontiers in Neuroinformatics, 2. https://doi.org/10.3389/neuro.11.010.2008

Pereverzeva, M., & Murray, S. O. (2008). Neural activity in human V1 correlates with dynamic lightness induction. Journal of Vision, 8(15), 8.1-10. https://doi.org/10.1167/8.15.8

Perna, A., Tosetti, M., Montanaro, D., & Morrone, M. C. (2005). *Neuron*al mechanisms for illusory brightness perception in humans. Neuron, 47(5), 645–651. https://doi.org/10.1016/j.neuron.2005.07.012

Pessoa, L., Mingolla, E., & Neumann, H. (1995). A contrast- and luminance-driven multiscale network model of brightness perception. Vision Research, 35(15), 2201–2223.

Pinheiro, J., Bates, D., DebRoy, S., Sarkar, D., & R Core Team. (2017). nlme: Linear and Nonlinear Mixed Effects Models. Retrieved from https://CRAN.R-project.org/package=nlme

Polimeni, J. R., Fischl, B., Greve, D. N., & Wald, L. L. (2010). Laminar analysis of 7T BOLD using an imposed spatial activation pattern in human V1. NeuroImage, 52(4), 1334–1346. https://doi.org/10.1016/j.neuroimage.2010.05.005

Poser, B. A., Koopmans, P. J., Witzel, T., Wald, L. L., & Barth, M. (2010). Three dimensional echo-planar imaging at 7 Tesla. NeuroImage, 51(1), 261–266. https://doi.org/10.1016/j.neuroimage.2010.01.108

R Core Team. (2017). R: A Language and Environment for Statistical Computing. Retrieved from https://www.R-project.org

Rao, R. P., & Ballard, D. H. (1999). Predictive coding in the visual cortex: a functional interpretation of some extra-classical receptive-field effects. Nature Neuroscience, 2(1), 79–87. https://doi.org/10.1038/4580

Ress, D., Glover, G. H., Liu, J., & Wandell, B. (2007). Laminar profiles of functional activity in the human brain. NeuroImage, 34(1), 74–84. https://doi.org/10.1016/j.neuroimage.2006.08.020

Rezak, M., & Benevento, L. A. (1979). A comparison of the organization of the projections of the dorsal lateral geniculate nucleus, the inferior pulvinar and adjacent lateral pulvinar to primary visual cortex (area 17) in the macaque monkey. Brain Research, 167(1), 19–40.

Rockland, K. S., & Pandya, D. N. (1979). Laminar origins and terminations of cortical connections of the occipital lobe in the rhesus monkey. Brain Research, 179(1), 3–20. https://doi.org/10.1016/0006-8993(79)90485-2

Rockland, K. S., & Virga, A. (1989). Terminal arbors of individual ‘feedback’ axons projecting from area V2 to V1 in the macaque monkey: a study using immunohistochemistry of anterogradely transported Phaseolus vulgaris-leucoagglutinin. The Journal of Comparative Neurology, 285(1), 54–72. https://doi.org/10.1002/cne.902850106

Roe, A. W., Lu, H. D., & Hung, C. P. (2005). Cortical processing of a brightness illusion. Proceedings of the National Academy of Sciences of the United States of America, 102(10), 3869–3874. https://doi.org/10.1073/pnas.0500097102

Roelfsema, P. R., Lamme, V. A. F., Spekreijse, H., & Bosch, H. (2002). Figure-ground segregation in a recurrent network architecture. Journal of Cognitive Neuroscience, 14(4), 525–537. https://doi.org/10.1162/08989290260045756

Rossi, A. F., & Paradiso, M. A. (1999). Neural correlates of perceived brightness in the retina, lateral geniculate nucleus, and striate cortex. The Journal of Neuroscience: The Official Journal of the Society for Neuroscience, 19(14), 6145–6156.

Rossi, A. F., Rittenhouse, C. D., & Paradiso, M. A. (1996). The representation of brightness in primary visual cortex. Science (New York, N.Y.), 273(5278), 1104–1107.

Saalmann, Y. B., Pinsk, M. A., Wang, L., Li, X., & Kastner, S. (2012). The pulvinar regulates information transmission between cortical areas based on attention demands. Science (New York, N.Y.), 337(6095), 753–756. https://doi.org/10.1126/science.1223082

Sasaki, Y., & Watanabe, T. (2004). The primary visual cortex fills in color. Proceedings of the National Academy of Sciences of the United States of America, 101(52), 18251–18256. https://doi.org/10.1073/pnas.0406293102

Schnabel, U. H., Kirchberger, L., van Beest, E., Mukherjee, S., Barsegyan, A., Lorteije, J. A. M., … Roelfsema, P. R. (2018). Feedforward and feedback processing during figure-ground perception in mice. BioRxiv. https://doi.org/10.1101/456459

Seghier, M., Dojat, M., Delon-Martin, C., Rubin, C., Warnking, J., Segebarth, C., & Bullier, J. (2000). Moving illusory contours activate primary visual cortex: an fMRI study. Cerebral Cortex (New York, N.Y.: 1991), 10(7), 663–670.

Self, M. W., van Kerkoerle, T., Supèr, H., & Roelfsema, P. R. (2013). Distinct roles of the cortical layers of area V1 in figure-ground segregation. Current Biology : CB, 23(21), 2121–2129. https://doi.org/10.1016/j.cub.2013.09.013

Setsompop, K., Gagoski, B. A., Polimeni, J. R., Witzel, T., Wedeen, V. J., & Wald, L. L. (2012). Blipped-controlled aliasing in parallel imaging for simultaneous multislice echo planar imaging with reduced g-factor penalty. Magnetic Resonance in Medicine, 67(5), 1210–1224. https://doi.org/10.1002/mrm.23097

Sherman, S. M., & Guillery, R. W. (2002). The role of the thalamus in the flow of information to the cortex. Philosophical Transactions of the Royal Society of London. Series B, Biological Sciences, 357(1428), 1695–1708. https://doi.org/10.1098/rstb.2002.1161

Shimono, M., Mano, H., & Niki, K. (2012). The brain structural hub of interhemispheric information integration for visual motion perception. Cerebral Cortex (New York, N.Y.: 1991), 22(2), 337–344. https://doi.org/10.1093/cercor/bhr108

Shmuel, A., Augath, M., Oeltermann, A., & Logothetis, N. K. (2006). Negative functional MRI response correlates with decreases in neuronal activity in monkey visual area V1. Nature Neuroscience, 9(4), 569–577. https://doi.org/10.1038/nn1675

Shmuel, A., Yacoub, E., Pfeuffer, J., Van de Moortele, P. F., Adriany, G., Hu, X., & Ugurbil, K. (2002). Sustained negative BOLD, blood flow and oxygen consumption response and its coupling to the positive response in the human brain. Neuron, 36(6), 1195–1210.

Smith, S. M., Jenkinson, M., Woolrich, M. W., Beckmann, C. F., Behrens, T. E. J., Johansen-Berg, H., … Matthews, P. M. (2004). Advances in functional and structural MR image analysis and implementation as FSL. NeuroImage, 23, S208–S219. https://doi.org/10.1016/j.neuroimage.2004.07.051

Standage, G. P., & Benevento, L. A. (1983). The organization of connections between the pulvinar and visual area MT in the macaque monkey. Brain Research, 262(2), 288–294.

Tosun, D., Rettmann, M. E., Han, X., Tao, X., Xu, C., Resnick, S. M., … Prince, J. L. (2004). Cortical surface segmentation and mapping. NeuroImage, 23, S108–S118. https://doi.org/10.1016/j.neuroimage.2004.07.042

Trojanowski, J. Q., & Jacobson, S. (1977). The morphology and laminar distribution of cortico-pulvinar neurons in the rhesus monkey. Experimental Brain Research, 28(1–2), 51–62.

Uludağ, K. (2008). Transient and sustained BOLD responses to sustained visual stimulation. Magnetic Resonance Imaging, 26(7), 863–869. https://doi.org/10.1016/j.mri.2008.01.049

Uludağ, K., & Blinder, P. (2018). Linking brain vascular physiology to hemodynamic response in ultra-high field MRI. NeuroImage, 168, 279–295. https://doi.org/10.1016/j.neuroimage.2017.02.063

van der Walt, S., Colbert, S. C., & Varoquaux, G. (2011). The NumPy Array: A Structure for Efficient Numerical Computation. Computing in Science & Engineering, 13(2), 22–30. https://doi.org/10.1109/MCSE.2011.37

Van Essen, D. C., Newsome, W. T., Maunsell, J. H., & Bixby, J. L. (1986). The projections from striate cortex (V1) to areas V2 and V3 in the macaque monkey: asymmetries, areal boundaries, and patchy connections. The Journal of Comparative Neurology, 244(4), 451–480. https://doi.org/10.1002/cne.902440405

van Kerkoerle, T., Self, M. W., Dagnino, B., Gariel-Mathis, M.-A., Poort, J., van der Togt, C., & Roelfsema, P. R. (2014). Alpha and gamma oscillations characterize feedback and feedforward processing in monkey visual cortex. Proceedings of the National Academy of Sciences, 111(40), 14332–14341. https://doi.org/10.1073/pnas.1402773111

Villeneuve, M. Y., Kupers, R., Gjedde, A., Ptito, M., & Casanova, C. (2005). Pattern-motion selectivity in the human pulvinar. NeuroImage, 28(2), 474–480. https://doi.org/10.1016/j.neuroimage.2005.06.015

Villeneuve, M. Y., Thompson, B., Hess, R. F., & Casanova, C. (2012). Pattern-motion selective responses in MT, MST and the pulvinar of humans. The European Journal of Neuroscience, 36(6), 2849–2858. https://doi.org/10.1111/j.1460-9568.2012.08205.x

von der Heydt, R., Friedman, H. S., & Zhou, H. (2003). Searching for the neural mechanism of color filling-in. In L. Pessoa & P. De Weerd, *Filling-in: from perceptual completion to cortical reorganization* (pp. 106–127). https://doi.org/10.1093/acprof:oso/9780195140132.001.0001

von der Heydt, R., Peterhans, E., & Baumgartner, G. (1984). Illusory contours and cortical neuron responses. Science (New York, N.Y.), 224(4654), 1260–1262.

Waehnert, M. D., Dinse, J., Weiss, M., Streicher, M. N., Waehnert, P., Geyer, S., … Bazin, P.-L. (2014). Anatomically motivated modeling of cortical laminae. NeuroImage, 93, 210–220. https://doi.org/10.1016/j.neuroimage.2013.03.078

Wokke, M. E., Vandenbroucke, A. R. E., Scholte, H. S., & Lamme, V. A. F. (2013). Confuse your illusion: feedback to early visual cortex contributes to perceptual completion. Psychological Science, 24(1), 63–71. https://doi.org/10.1177/0956797612449175

Wong-Riley, M. T. (1974). Demonstration of geniculocortical and callosal projection neurons in the squirrel monkey by means of retrograde axonal transport of horseradish peroxidase. Brain Research, 79(2), 267–272.

Yushkevich, P. A., Piven, J., Hazlett, H. C., Smith, R. G., Ho, S., Gee, J. C., & Gerig, G. (2006). User-guided 3D active contour segmentation of anatomical structures: Significantly improved efficiency and reliability. NeuroImage, 31(3), 1116–1128. https://doi.org/10.1016/j.neuroimage.2006.01.015

Zhang, Y., Brady, M., & Smith, S. (2001). Segmentation of brain MR images through a hidden Markov random field model and the expectation-maximization algorithm. IEEE Transactions on Medical Imaging, 20(1), 45–57. https://doi.org/10.1109/42.906424

Zhao, F., Wang, P., & Kim, S.-G. (2004). Cortical depth-dependent gradient-echo and spin-echo BOLD fMRI at 9.4T. Magnetic Resonance in Medicine, 51(3), 518–524. https://doi.org/10.1002/mrm.10720

Zipser, K., Lamme, V. A., & Schiller, P. H. (1996). Contextual modulation in primary visual cortex. The Journal of Neuroscience: The Official Journal of the Society for Neuroscience, 16(22), 7376–7389.

