## Supplementary Material for "Depth-resolved ultra-high field fMRI reveals feedback contributions to surface motion perception"

### Population receptive field mapping

Stimuli used for population receptive field mapping were oriented bars at four different orientations and eight different positions per orientation, containing a black and white checkerboard pattern. The bars had a width of 1.25° visual angle, and the carrier pattern within the bar had a spatial frequency of 1.2 cycles/deg. The luminance of the black and white sectors of the carrier pattern was 2 cd/m2 and 1390 cd/m2, respectively, resulting in a luminance contrast of 1. The polarity of the checkerboard pattern was reversed at a frequency of 4 Hz, and the bar changed its position every 2.079 s (in synchrony with the volume TR). Each of the resulting 32 stimulus configurations was presented 12 times for 2.08 s in random order. The duration of the population receptive field mapping run was 832 s (400 volumes). Similar to the main experiment, subjects were instructed to perform a central fixation task during the retinotopic mapping experiment. The software used for the presentation of the retinotopic mapping stimuli is publicly available (https://doi.org/10.5281/zenodo.1475439).

### Supplementary Figures

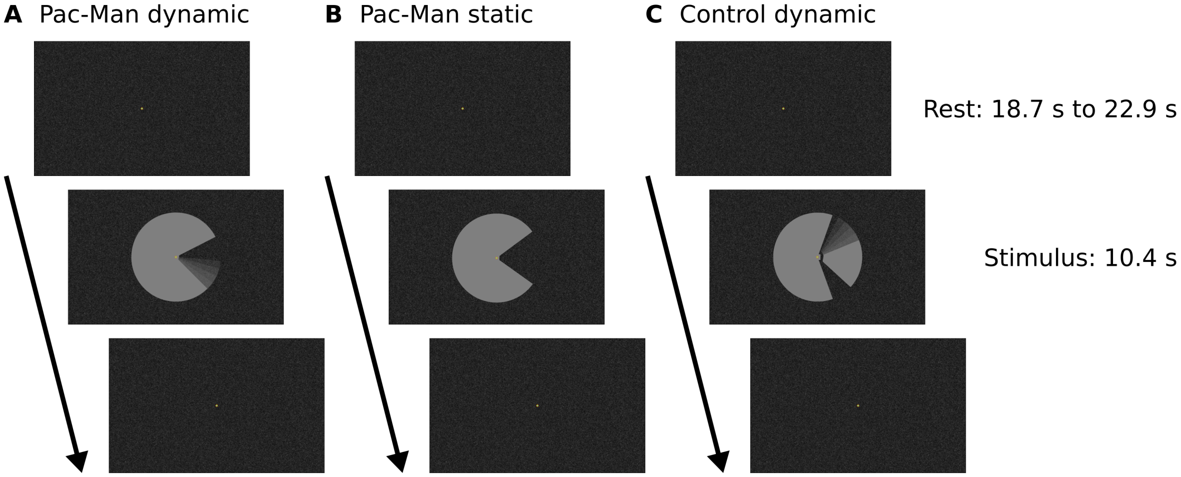


*Supplementary Figure S1.* Experimental design. Stimuli were presented in a block design with rest blocks of variable duration. The three stimulus conditions were presented in separate runs (A, B, C). A central fixation dot and the random texture background pattern were present throughout the duration of each run.

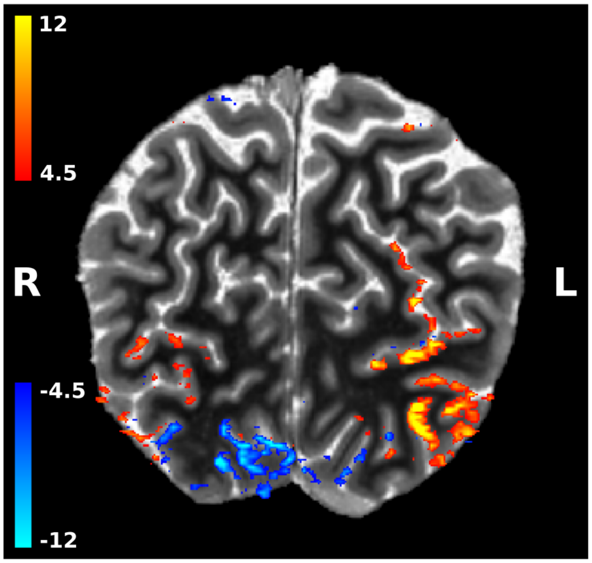


*Supplementary Figure S2.* The Pac-Man stimulus caused positive and negative fMRI signal changes across visual cortex. Shown are the z-scores for the GLM contrast Pac-Man dynamic (sustained response) against rest, overlaid on a brain-masked T1 image, for a representative subject. Negative signal changes are particularly pronounced in early visual cortex of the right hemisphere, i.e. the hemisphere that 'sees' the left side of the Pac-Man. (Image is in radiological convention.)

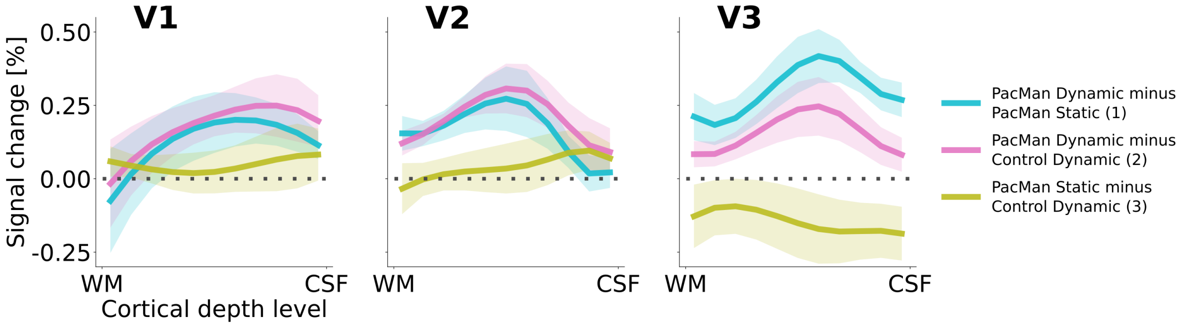


*Supplementary Figure S3.* Cortical depth profiles of condition differences. There are three possible condition contrasts: (1) PacMan dynamic vs. PacMan static (blue line), (2) PacMan dynamic vs. control dynamic (magenta line), and (3) PacMan static vs. control dynamic (yellow line). Shading represents the standard error of the mean (across subjects). The spatial deconvolution for removal of signal spread due to draining veins was applied to the individual condition depth profiles separately for each subject.

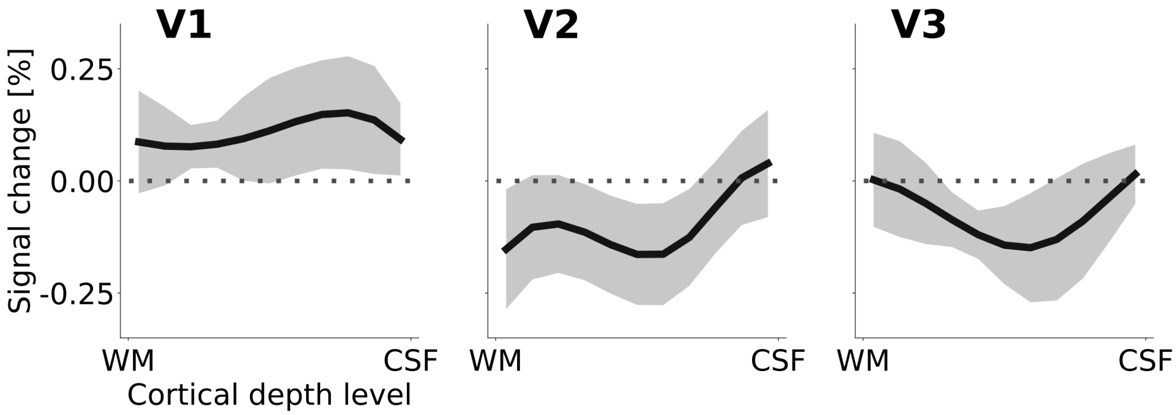


*Supplementary Figure S4.* Cortical depth profiles of the apparent motion effect for the cortical representation of the stimulus edge (see Figure 2 C & F). The apparent motion effect was defined as the relative signal change associated with the condition contrast 'Pac-Man dynamic' (Figure 1 A) minus 'control dynamic' (Figure 1 C). Shading represents the standard error of the mean (across subjects).

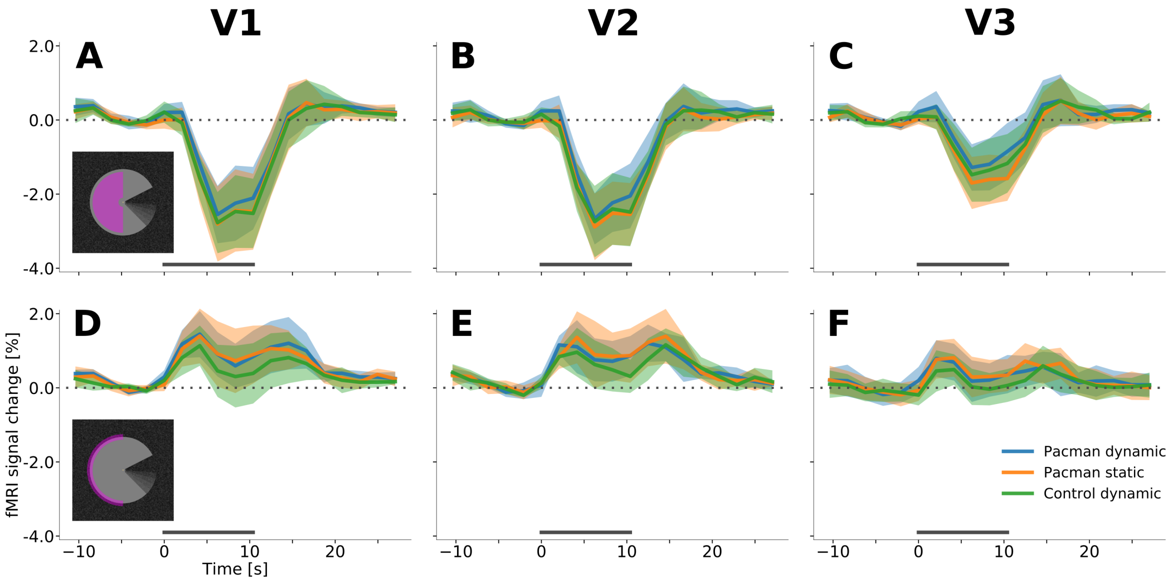


*Supplementary Figure S5.* Event-related fMRI timecourses for region of interest corresponding to the stimulus centre (A, B, C) and the edge of the stimulus (D, E, F) in the right hemisphere. The horizontal grey bar marks the duration of the stimulus block. All three stimulus conditions (represented by separate lines) evoked a sustained negative response in V1, V2, and V3 in cortex that retinotopically represents the stimulus centre. In contrast, the cortex that represents the stimulus edge exhibits a transient, positive response at stimulus onset and stimulus offset. Interestingly, the positive response at the stimulus edge precedes the negative response at the stimulus centre (see also Figure 4 in the main text). Error shading represents the standard error of the mean (across subjects). The scale of the axes is identical in all subplots.

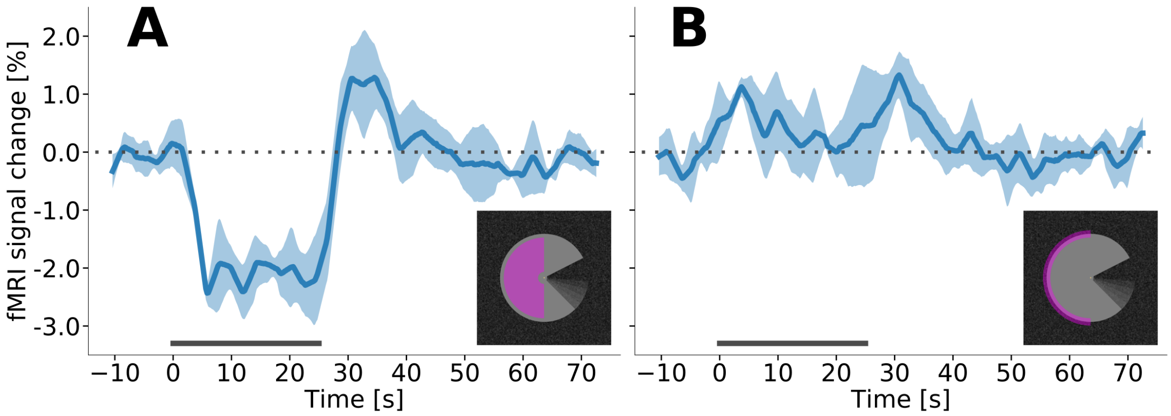


*Supplementary Figure S6.* Event-related fMRI timecourses from an additional run with longer stimulus blocks, for V1 in the right hemisphere. In order to further investigate the temporal dynamics of the stimulus-evoked response, we acquired and additional run during which the dynamic Pac-Man stimulus was presented with longer block durations (in a subset of subjects, $n$=5). Stimulus duration was 25 s, with rest blocks of 50 s. (A) The region of interest that corresponds to the stimulus centre shows a sustained, negative response that resembles the 'canonical' haemodynamic response function. (B) The response to the stimulus edge is transient and positive.

### Supplementary Table

*Supplementary Table 1.* Comparison of stimulus parameters in Akin et al. (2014) and in the present study. Akin, B., Ozdem, C., Eroglu, S., Keskin, D. T., Fang, F., Doerschner, K., ... Boyaci, H. (2014). Attention modulates neuronal correlates of interhemispheric integration and global motion perception. *Journal of Vision*, 14(12). https://doi.org/10.1167/14.12.30

| | **Akin et al. 2014** | **Present study** |
|---|---|---|
| Diameter of stimulus | 15° visual angle | 7.5° visual angle |
| Viewing mode | Central fixation task & passive viewing | Central fixation task |
| Rest block duration | 12 s | 18.7 s, 20.8 s, or 22.9 s |
| Stimulus block duration | 12 s | 10.4 s |
| Stimulus luminance | 503 cd/m$^2$ | 163 cd/m$^2$ |
| Mean background luminance | 189 cd/m$^2$ | 8 cd/m$^2$ |
| Oscillation rate of stimulus | 1.04 Hz | 0.85 Hz |
